# Substrate stiffness dictates unique paths towards proliferative arrest in WI-38 cells

**DOI:** 10.1101/2024.09.24.614744

**Authors:** Alyssa M. Kaiser, Amirali Selahi, Wenjun Kong, J. Graham Ruby

**Affiliations:** Calico Life Sciences LLC, South San Francisco, United States

## Abstract

Finite replicative potential is a defining feature of non-transformed somatic cells, first established by Leonard Hayflick *in vitro* using WI-38 human lung fibroblasts. Once proliferative capacity is exhausted due to telomere shortening, cells enter into a state called replicative senescence, which can be avoided through ectopic expression of telomerase reverse transcriptase (hTERT). As WI-38 cells approach replicative arrest, molecular pathways linked to mechanotransduction are induced, including YAP signaling, but the potential interplay between replicative lifespan and the mechanical environment of the cell remains unexplored. Here, we investigate the influence of mechanosensation on the trajectory towards replicative arrest taken by WI-38 cells by growing cells on substrates of varying stiffnesses. Matrix softening slowed proliferation, altered cellular phenotypes, and shortened proliferative lifespan while hTERT expression abrogated or reduced these responses. Our analyses of bulk and single-cell RNA-sequencing and ATAC-sequencing revealed the emergence of a unique G1 transcriptional state on soft substrates, characterized by an AP-1 transcription factor program, which failed to manifest with hTERT expression. Together, these findings reveal how the mechanical environment alters WI-38 cell proliferative lifespan and dictates unique paths towards growth arrest.

## Introduction

The phenomenon that primary mammalian cells undergo a finite number of population doublings before undergoing replicative senescence was first discovered by Leonard Hayflick in 1965^1^. Since then, telomere erosion has been identified as the major causal process driving replicative senescence, triggering a DNA damage response and growth arrest once telomeres are insufficiently long to protect the ends of chromosomes^2–4^. Expression of telomerase reverse transcriptase (hTERT) allows cells to extend telomeres and is sufficient for indefinite proliferation, demonstrating the importance of telomere erosion in triggering replicative senescence^3^. Other interventions that extend replicative lifespan target cellular stresses and their response pathways, though they stop short of immortalizing cells. For instance, treating cells with antioxidants has been shown to increase the number of population doublings of human fibroblasts^5^. Additionally, inhibiting mTOR with rapamycin increased the replicative lifespan of bone marrow mesenchymal stem cells^6^. However, other potential modifiers of replicative lifespan remain to be explored, including perturbations to the mechanical properties of the physical environment.

Modern studies have dissected transcriptomic, metabolomic, epigenomic, secretory, and other intracellular changes that occur as cells age towards replicative senescence^7–9^. In their multiomics dissection of replicative lifespan, Chan et al.^7^ identified a potential connection between the extracellular environment and replicative lifespan. As WI-38 human fetal lung fibroblasts age, they undergo a fibroblast-to-myofibroblast (FMT) transition, with a concomitant induction of myofibroblast identity, YAP target genes, and extracellular matrix (ECM) gene expression^7^. In contrast, cells experiencing irradiation-induced senescence or growth arrest via contact inhibition do not undergo FMT, emphasizing that WI-38 cells can follow distinct trajectories to proliferative arrest. A central theme among the changes associated with replicative senescence induction is the connection to mechanotransduction and the mechanical environment, as extracellular forces can promote YAP activation and induce myofibroblast identity^10,11^. Treating late passage WI-38 cells with the YAP inhibitor verteporfin reversed FMT, YAP, and ECM programs^7^. These data identifying YAP, a mediator of mechanosensory signals^10^, as a potential regulator of replicative senescence phenotypes suggest a relationship between cellular senescence and the mechanical environment.

An important aspect of the mechanical environment is its elastic modulus, or stiffness, a variable relevant to the aging process: many tissues stiffen with age, including cardiovascular^12^ and pulmonary^13^ tissues. Additionally, tissue stiffening can contribute to disease and aging phenotypes. For example, aortic stiffening contributes to structural abnormalities in the heart, potentially resulting in heart failure^12^. Understanding how this variable contributes to the aging process *in vitro* may provide further insights towards more complex systems. As WI-38 cells were intolerant to long-term culturing with verteporfin (see Discussion), could dampening YAP activity and FMT pathways via alternative mechanisms, such as a reduction in environmental stiffness, extend replicative lifespan?

We sought to experimentally test the hypothesis that reducing the mechanical force exerted through the extracellular environment would reverse transcriptional programs associated with replicative senescence and increase the number of population doublings undergone by WI-38 cells. To manipulate the mechanical force experienced by cells, we altered the stiffness of the substrates cells are cultured on, as decreasing matrix stiffness reduces external force on cells^14^. Here, we perform the first longitudinal study of how substrate stiffness affects replicative lifespan and investigate how environmental stiffness affects proliferative, transcriptional, epigenomic, and other cellular phenotypes over time. We discover that matrix softening shortens the proliferative lifespan of wild-type WI-38 cells, concomitant with the emergence of a unique G1 transcriptional state and distinct transcription factor programs, while hTERT expression abrogates many of the phenotypes induced by matrix softening.

## Results

### Matrix stiffness influences WI-38 cell proliferation rate and capacity

To understand how matrix stiffness affects WI-38 cell biology and replicative lifespan, we designed a time course experiment to longitudinally cultivate and sample cells on different substrate stiffnesses. These substrates encompassed a wide range of elastic moduli, the measurement of a material’s elasticity. Cells were grown on a “stiff” substrate, collagen-coated plastic tissue culture plates that have an elastic modulus of 2.3 gigapascals (GPa)^15,16^. To contrast a stiff extracellular environment, cells were grown on “soft” substrates - collagen-coated polydimethylsiloxane (PDMS) gels ranging from 0.5 to 32 kilopascals (kPa). This range mimics the physiological stiffness of healthy (0.5 and 2 kPa) and fibrotic (16 and 32 kPa) lungs^17^. Early passage (population doubling level 22, PDL22) wild-type WI-38 cells were grown on the stiff and soft surfaces for approximately five months and periodically collected for gene expression analysis and senescence-associated β-galactosidase (SA-β-gal) staining (Fig. 1a,b). WI-38 cells immortalized with hTERT were grown in parallel as a control for cells that would not undergo replicative senescence.

**Figure 1:**
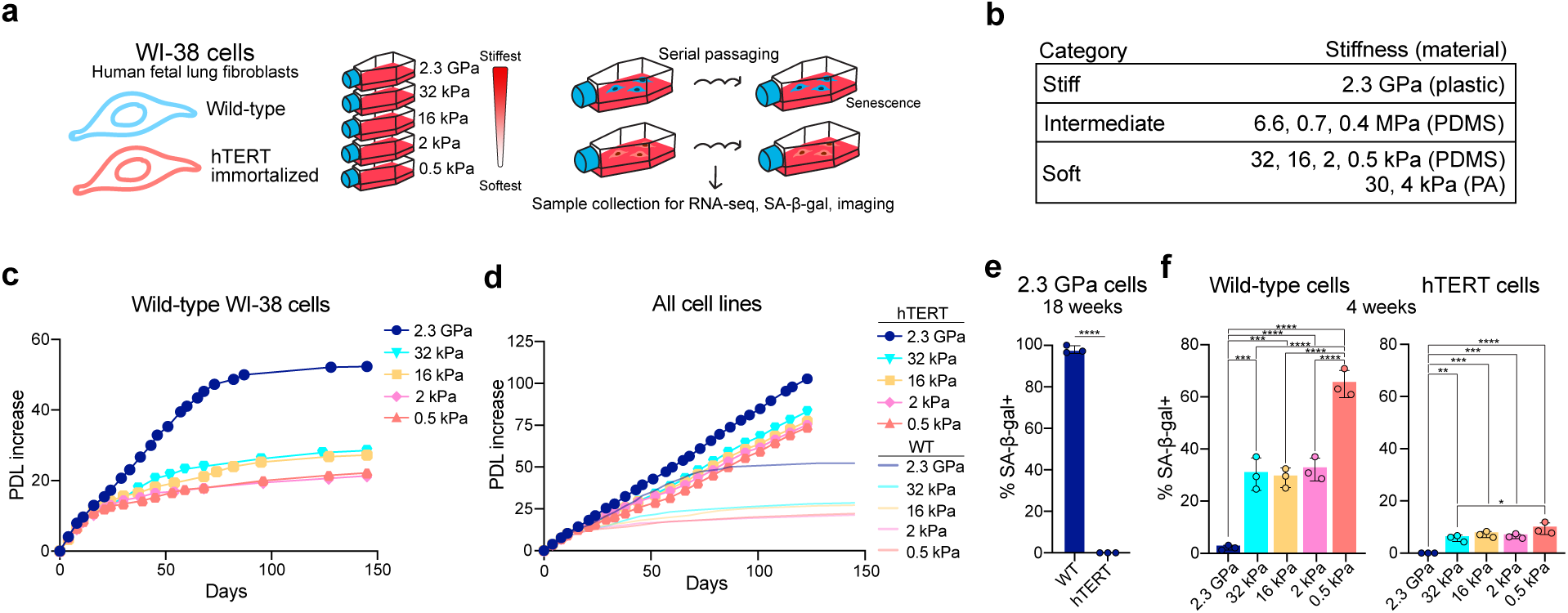
Matrix stiffness affects WI-38 cell proliferation rate. (a) Schematic for the time course experiment. GPa, gigapascal; kPa, kilopascal. (b) Elastic moduli (stiffness) ranges of stiff, intermediate, and soft substrates. PDMS, polydimethylsiloxane; PA, polyacrylamide; MPa, megapascal. (c) Line graph displaying days in culture (x-axis) versus the population doubling level (PDL) increase from day 0 (y-axis) for wild-type WI-38 cells on different matrices. (d) Line graph displaying the days in culture (x-axis) versus the PDL increase from day 0 (y-axis) for wild-type (WT) and hTERT-expressing WI-38 cells on different matrices. (e) Bar graph quantifying the percent of cells that stained for senescence-associated β-galactosidase (SA-β-gal) in wild-type and hTERT-expressing WI-38 cells after 18 weeks in culture. (f) Bar graphs quantifying the percent of cells that stained for SA-β-gal in (left) wild-type and (right) hTERT-expressing WI-38 cells after 4 weeks in culture across substrates. Bar graphs are mean ± standard deviation. *P* values were calculated by two-tailed Student’s *t*-test (e) or ordinary one-way ANOVA with Tukey’s multiple comparisons test (f). \**P* < 0.05, \*\**P* < 0.01, \*\*\**P* < 0.001, \*\*\*\**P* < 0.0001.

Wild-type WI-38 cells grew robustly on the stiff surface for just over three months before undergoing an abrupt growth arrest, consistent with previous reports (Fig. 1c)^7^. Over the first month, wild-type cells across all stiffnesses reached similar population doubling levels (Fig. 1c). After this time, WI-38 cells showed a stiffness-dependent change in proliferation; the number of population doublings underwent decreased with matrix softening (Fig. 1c). Thus, matrix softening reduces the proliferative lifespan of wild-type WI-38 cells. Meanwhile, hTERT-immortalized WI-38 cells on the stiff substrate exhibited a consistent growth rate throughout the entire experiment, as expected (Fig. 1d)^7^. hTERT cells on the soft matrices initially displayed a modest reduction in proliferation rate, but thereafter grew at a constant rate over the remainder of the experiment (Fig. 1d).

While matrix softening correlated with fewer population doublings achieved before wild-type cells underwent a growth arrest, it is possible that this was due to a differential response to plastic versus PDMS surfaces. To address this concern, we leveraged two distinct approaches. First, we fabricated PDMS plates^18–20^ of intermediate stiffnesses, multiple orders of magnitude stiffer than the soft matrices previously used, to test how growth on an intermediate stiffness affected the number of doublings undergone by WI-38 cells (Fig. S1a, Supplemental Data File 1). Wild-type (PDL29) and hTERT WI-38 cells were grown on stiff (2.3 GPa; plastic), intermediate (6.6, 0.7, 0.4 megapascals [MPa]; PDMS), and soft (2 kPa; PDMS) collagen-coated substrates until the wild-type cells ceased proliferating. Wild-type cells on the soft matrix underwent a rapid proliferative arrest after 3-4 weeks in culture while those on the stiff substrate proliferated for over 10 weeks, consistent with our initial observations (Fig. S1b). Wild-type cells grown on the intermediate stiffnesses reached an intermediate PDL between the soft and stiff plates before ultimately arresting (Fig. S1b). These data suggest matrix softening, rather than surface material, reduces the proliferative lifespan of wild-type WI-38 cells.

To further exclude the possibility that our findings were the result of a specific response to PDMS, we additionally cultured cells on soft polyacrylamide (PA) matrices to test if phenotypes were consistent across distinct soft materials. Wild-type (PDL29) and hTERT WI-38 cells were grown on stiff (2.3 GPa; plastic), soft PA (4 kPa and 30 kPa), and soft PDMS (2 kPa) collagen-coated substrates (Fig. S1c). Wild-type WI-38 cells on 2 kPa PDMS and 4 kPa PA matrices exhibited nearly identical growth, undergoing a rapid growth arrest after 4 weeks of culturing, while cells grown on 30 kPa PA proliferated to a slightly greater PDL (Fig. S1d). As expected, cells on the stiff substrate grew consistently throughout this time frame (Fig. S1d). Thus, matrix softening reduces the proliferative lifespan of wild-type WI-38 cells across a range of elastic moduli and substrates.

### Substrate softening induces senescence-associated phenotypes prematurely in wild-type WI-38 cells

Throughout the experiment, WI-38 cells were examined for features of senescence. Cells were collected periodically for SA-β-bal staining, an established marker of senescent cells^21^. Wild-type WI-38 cells grown on the stiff surface showed little SA-β-gal staining at early points in the time course, but staining drastically increased as the cells became nonproliferative (Fig. 1e,f, S2a,b). Only rare hTERT-expressing cells grown on the stiff substrate displayed SA-β-bal staining at any point in the time course (Fig. 1e,f, S2a,b). In contrast, SA-β-gal staining was observed in both wild-type and hTERT WI-38 cells grown on soft matrices as soon as after two weeks of growth, with increased staining in wild-type cells (Fig. S2a). SA-β-gal activity increased over time in cells on soft surfaces; after 4 weeks of growth, 30-60% of wild-type cells were SA-β-gal+ as were about 10% of hTERT cells (Fig. 1f, S2a).

In addition to earlier SA-β-gal induction, matrix softening expedited morphological changes associated with senescence in wild-type cells. Late passage wild-type cells grown on stiff plastic became enlarged and spread out as they ceased proliferating (Fig. S2b,c). On soft substrates, similar cell enlargement and spreading occurred as early as after three weeks of culturing (Fig. S2b,c). Across all stiffnesses, hTERT-expressing cells did not undergo visible changes to cell shape and/or size over time (Fig. S2b,c).

Wild-type WI-38 cells cultured on soft substrates exhibited proliferative, morphological, and SA-β-bal phenotypes within 20 population doublings that took over 50 population doublings to manifest in cells on stiff substrates. Across all surfaces and experiments, hTERT expression blocked the emergence of these phenotypes, even at early time points on soft substrates. In the case of the stiff substrate, these phenotypes are known to be coincident with replicative senescence^1,7^. In contrast, the premature manifestation of these phenotypes on soft matrices called into question whether cells on soft substrates underwent telomere-driven replicative senescence versus some other form of premature growth arrest.

### WI-38 cells display broadly similar senescence-associated transcriptional profiles across all surfaces

To understand how matrix softening reduces the proliferative lifespan of WI-38 cells, we performed RNA-sequencing on samples collected throughout the initial time course (Fig. 2a). Hierarchical clustering and principal component analysis of all samples revealed genotype and time point as the top contributors to transcriptional variance, with matrix stiffness accounting for a smaller percent of variance (Fig. 2b, S3a). As expected, given its established role in mechanosensation^10^, YAP pathway signaling in particular was rapidly reduced following initial plating of WI-38 cells onto soft matrices (Fig. S3b). We then examined the dataset to compare senescence-associated transcriptional phenotypes in wild-type WI-38 cells on stiff and soft matrices. Genes commonly induced in senescence, such as *CDKN1A*, *CDKN2A*, and *CDKN2B*^9^, were activated in wild-type cells on both stiff and soft surfaces over time (Fig. 2c).

**Figure 2:**
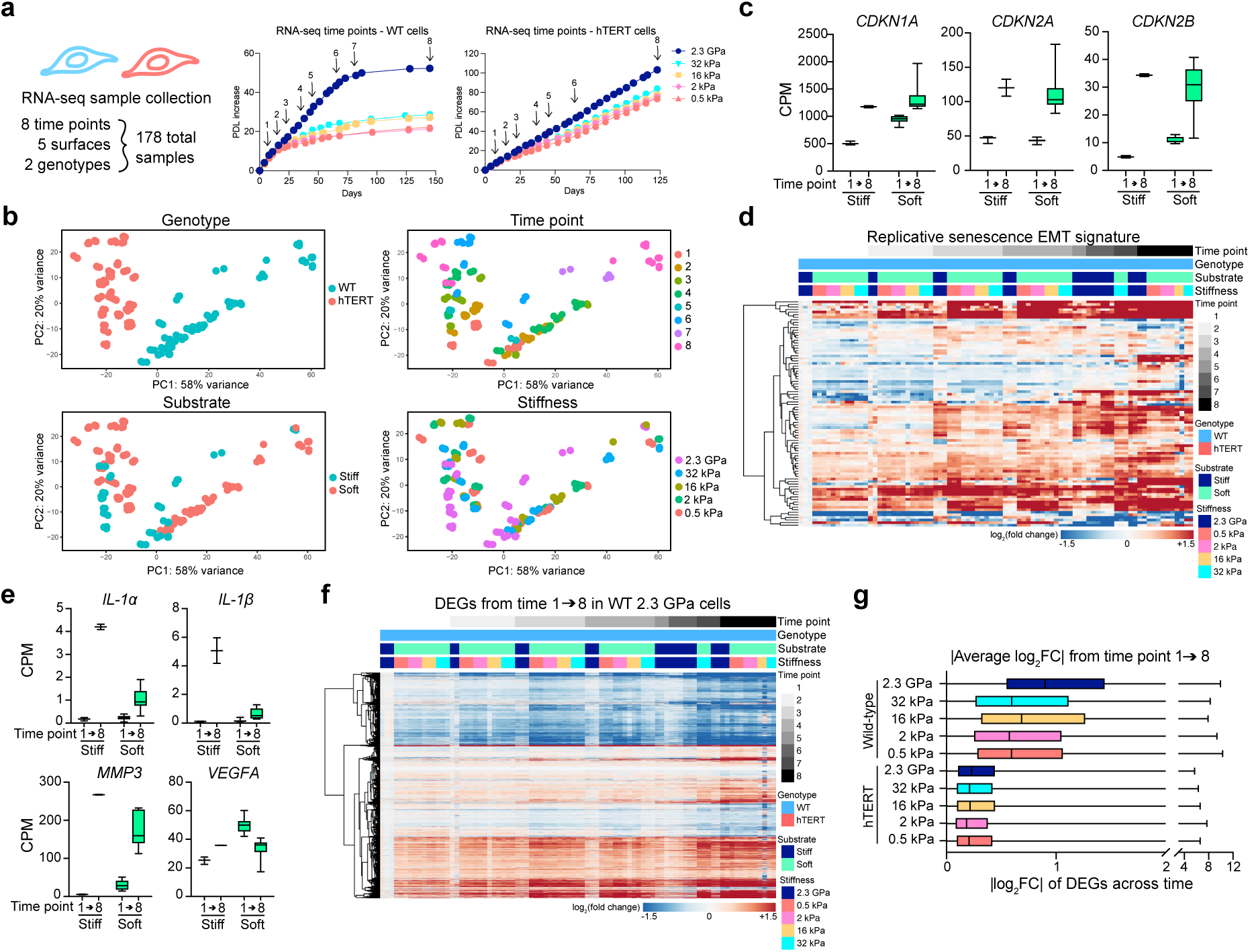
Wild-type WI-38 cells cultured on stiff and soft matrices similarly regulate senescence-associated gene expression programs. (a) (Left) Schematic of the RNA-sequencing samples and (right) sample collection time points. (b) Principal component (PC) analysis showing PC1 and PC2 with samples colored by different variables. (c) Box plots comparing the counts per million (CPM) of the indicated genes in wild-type WI-38 cells on the indicated surfaces at time points 1 and 8. (d) Heatmap of the log_2_(fold change) of the epithelial-to-mesenchymal (EMT) gene set (Supplemental Data File 2) previously identified^7^ as upregulated in replicative senescence. (e) Box plots comparing the expression of senescence-associated secretory phenotype (SASP) genes^22^ in wild-type WI-38 cells on the indicated surfaces at time points 1 and 8. (f) Heatmap of the log_2_(fold change) of all differentially expressed genes (DEGs) in wild-type cells grown on the 2.3 GPa substrate from time point 1 to time point 8 plotted across all wild-type samples across time. (g) Box plots quantifying the absolute value of the log_2_(fold change) of each gene in (f) from time point 1 to time point 8 averaged across replicates. Box plots show median ± 25th and 75th percentiles. Whiskers show minimum to maximum.

Previous work identified that cells undergoing replicative senescence, and not premature senescence induced by irradiation or growth arrest due to contact inhibition, specifically induce an epithelial-to-mesenchymal (EMT) gene signature^7^. This EMT gene set was upregulated over time in wild-type cells on the stiff substrate that underwent replicative senescence (Fig. 2d, S3c). Despite undergoing a premature proliferative arrest, wild-type cells on soft matrices showed an upregulation of this EMT signature over time as well (Fig. 2d, S3c, Supplemental Data File 2). Additional RNA-sequencing of wild-type cells grown on the intermediate stiffness substrates (6.6, 0.7, and 0.4 MPa) revealed a similar induction of the EMT signature over time (Fig. S4a-c). Likewise, additional gene signatures associated with replicative senescence^7^, including myofibroblast markers and YAP pathway target genes, were induced over the lifespan of the cell regardless of matrix stiffness (Fig. S3d-g). In contrast, markers of the senescence-associated secretory phenotype (SASP)^22^ displayed dissimilar regulation between stiff and soft substrates. Growth on soft substrates dampened the induction of SASP markers, such as *IL-1α* and *MMP3* (Fig. 2e, S3h).

Next, we defined the senescence-associated transcriptional program in wild-type cells on the stiff plastic substrate as genes differentially expressed between the first and final time points. Consistent with previous work^7^, the vast majority of gene expression changes in this program emerged in wild-type cells on the stiff substrate months before the induction of replicative senescence and intensified over time (Fig. 3f, S3i). This program was similarly regulated in wild-type cells on soft matrices in terms of the directionality and magnitude of induction relative to the first time point (Fig. 3g, S3j). In contrast, this senescence-associated program was largely unchanged in hTERT-expressing cells across stiffnesses, demonstrating that hTERT expression was sufficient to suppress transcriptional changes associated with replicative senescence induction (Fig. 3g, S3k). Likewise, the senescence-associated program in wild-type cells grown on soft and intermediate matrices was similarly regulated across time in wild-type cells on the stiff substrate (Fig. S3l, S4d-f). Thus, WI-38 cells cultured on a range of matrix stiffnesses exhibit overall similar transcriptional changes across their proliferative lifespans, including the upregulation of many gene expression programs associated with replicative senescence.

**Figure 3:**
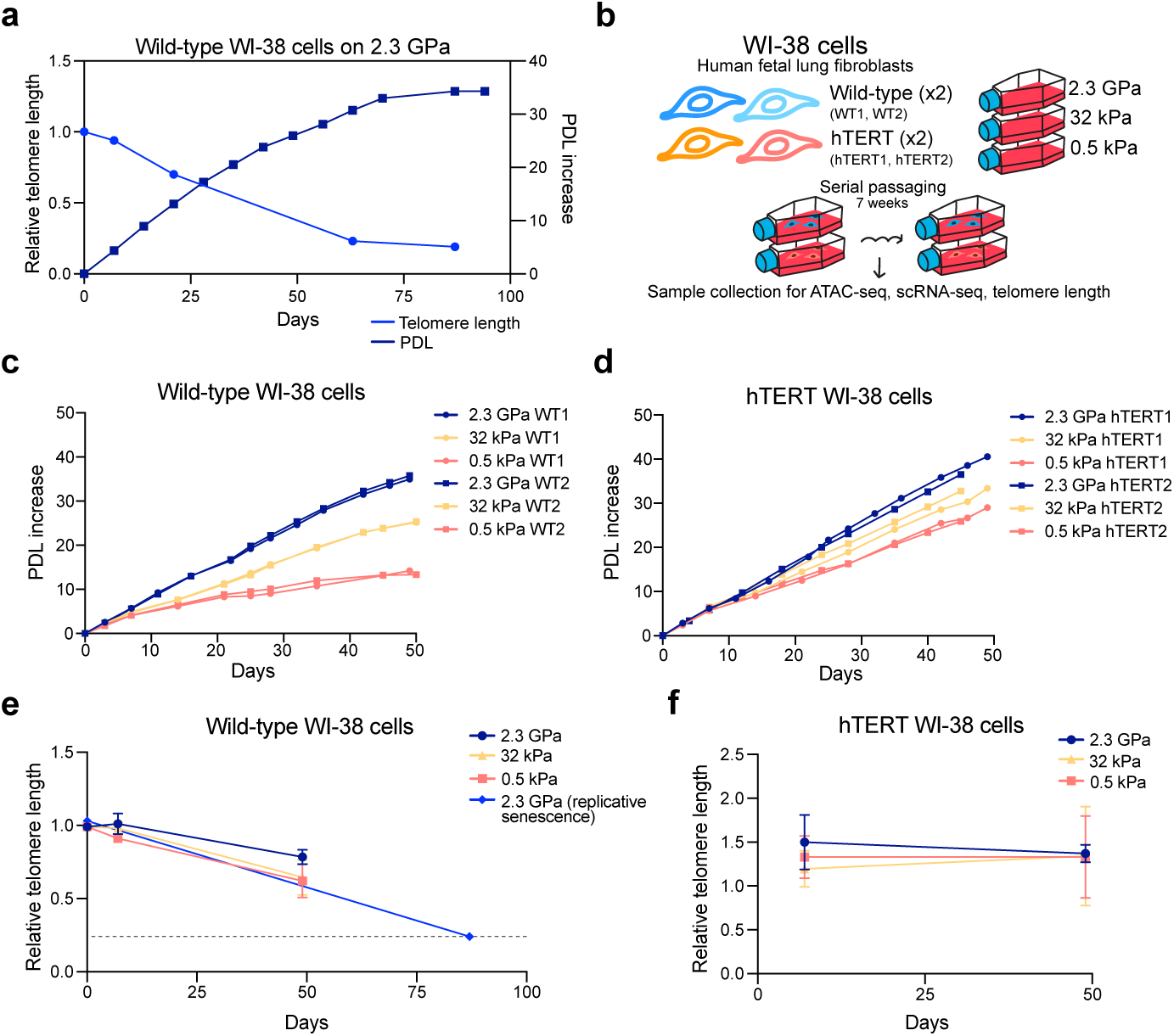
Telomeres are not critically shortened with matrix softening. (a) Line graph displaying days in culture (x-axis) versus relative telomere length (left y-axis) or versus the PDL increase (right y-axis) for wild-type WI-38 cells grown on stiff plastic. Samples are from the time course in Fig. S1b. (b) Schematic for the 7 week time course experiment. (c,d) Line graphs displaying the days in culture (x-axis) versus the PDL increase from day 0 (y-axis) for (c) wild-type and (d) hTERT-expressing WI-38 cells on different matrices. (e,f) Line graphs displaying days in culture (x-axis) versus the relative telomere length (y-axis) for (e) wild-type or (f) hTERT WI-38 cells on the indicated surfaces. The “2.3 GPa (replicative senescence) sample” is the same as in (a).

### Average telomere length is not critically shortened with matrix softening

Periodically through the replicative lifespan of wild-type cells grown on plastic, genomic DNA samples were collected for quantification of relative telomere lengths using quantitative PCR (qPCR)^23–25^. Overlapping relative telomere lengths with the growth curve revealed that telomeres were ∼80% shorter at the onset of replicative senescence than at time 0 (PDL29) (Fig. 3a). Given the earlier onset of replicative slowing and arrest and senescence-like phenotypes for wild-type WI-38 cells grown on soft substrates, we used qPCR to evaluate two alternative hypotheses: 1) that cells on soft substrates exhibited senescence-like behavior prior to critical telomere shortening, or 2) that the soft-substrate environment accelerated telomere shortening to induce replicative senescence earlier.

In order to collect samples for telomere quantification across a wider variety of conditions and acquire data from additional modalities (ATAC-sequencing and single-cell [sc]RNA-sequencing), we performed an additional time course experiment with wild-type and hTERT-expressing WI-38 cells (Fig. 3b). We used a sparser set of stiffnesses (2.3 GPa plastic, 0.5 kPa PDMS, 32 kPa PDMS) and limited to the first seven weeks of growth, a time period in which substrate-dependent phenotypes emerged in our initial experiment. Growth dynamics were reproduced here: the number of population doublings undergone by wild-type WI-38 cells decreased with matrix softening (Fig. 3c,d), and by day 50, wild-type cells on the 0.5 kPa surface ceased proliferating while those on the 32 kPa surface slowed but did not stop dividing (Fig. 3c).

Telomere qPCR was performed on samples harvested at 0, 1, and 7 weeks to measure relative telomere lengths^23–25^. Telomeres in wild-type WI-38 cells shortened over this time frame, with softer matrices modestly inducing greater telomere loss (Fig. 3e). Nonetheless, average decreases of 35% and 38% on the 32 kPa and 0.5 kPa surfaces, respectively, were far less than the ∼80% loss that coincided with replicative senescence on 2.3 GPa substrate (Fig. 3a,e). Meanwhile, over the 7 week time course, hTERT cells on all surfaces showed a consistent telomere length (Fig. 3f). Thus, while matrix softening accelerated the rate of telomere shortening, the level of degradation observed appeared insufficient for induction of replicative senescence.

### Matrix softening induces opening of AP-1 transcription factor binding sites

We next used ATAC-sequencing data to understand how matrix stiffness affects the epigenomic landscape. Samples were collected after 1, 4, and 7 weeks of culturing, and library quality was verified using fragment size distribution and enrichment around transcriptional start sites (Fig. 4a, S5a-d). Dimensionality reduction was performed using UMAP (uniform manifold approximation and projection), and akin to the bulk RNA-sequencing dataset, genotype was the main contributor to variance among the samples (Fig. 4b). Samples distribution was also dependent on time point and matrix stiffness, indicating that all three variables contributed to chromatin accessibility changes (Fig. 4b).

**Figure 4:**
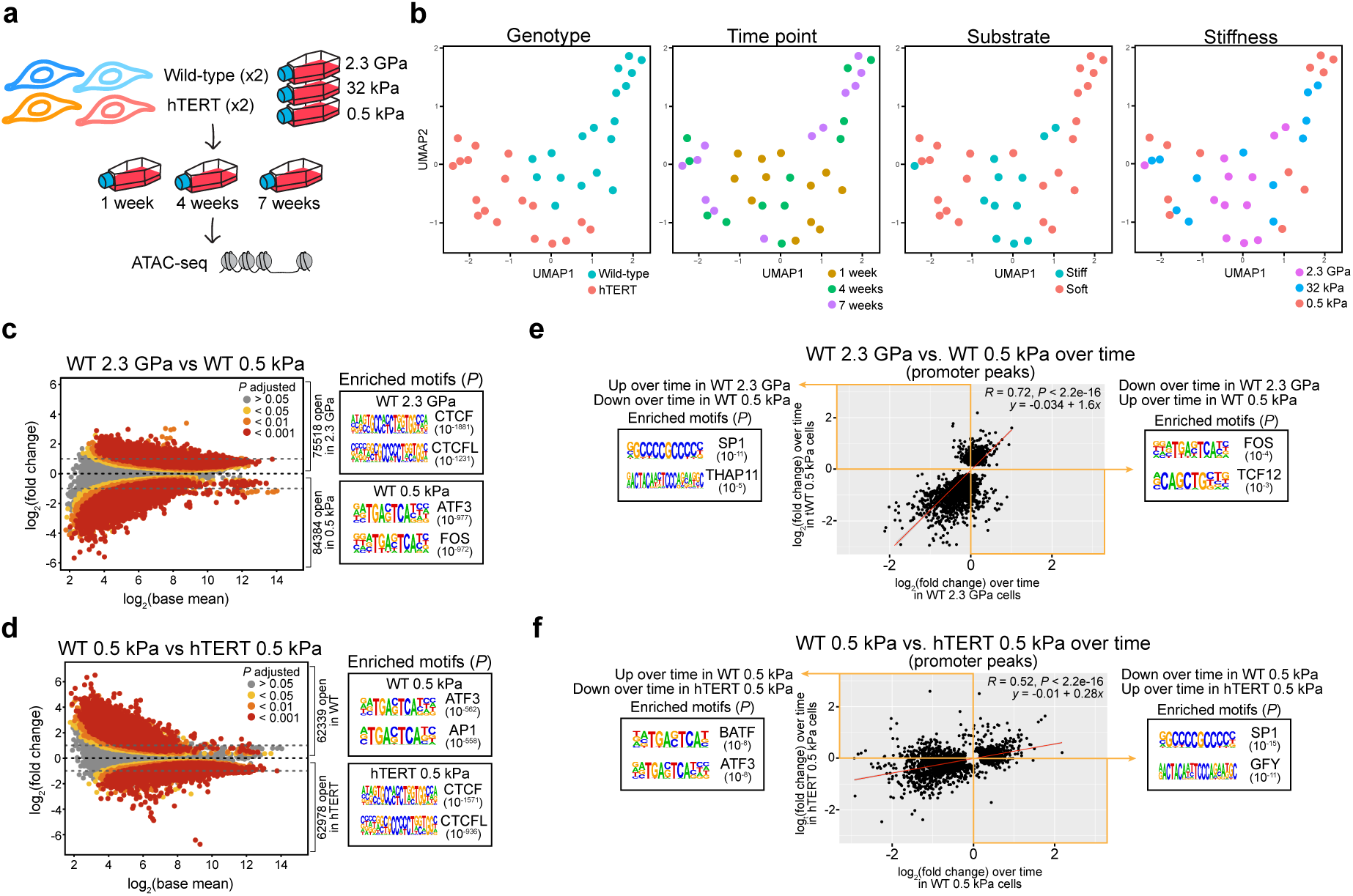
Matrix softening triggers opening of AP-1 transcription factor binding sites. (a) Schematic for the ATAC-sequencing experiment. (b) Uniform manifold approximation and projection (UMAP) projections of ATAC samples (all peaks analysis) showing UMAP1 and UMAP2 colored by different variables. (c,d) Log_2_ mean reads per region versus differential accessibility [log_2_(fold change)] for all peaks in the (c) wild-type 2.3 GPa versus wild-type 0.5 kPa comparison or the (d) wild-type 0.5 kPa versus hTERT 0.5 kPa comparison. The top 2 transcription factor motifs enriched in differentially accessible regions identified by HOMER known motif analysis are shown. (e,f) All differentially accessible promoter regions over time (time 1 versus time 7) in (e) wild-type cells on 2.3 GPa versus wild-type cells on 0.5 kPa or (f) wild-type cells on 0.5 kPa versus hTERT cells on 0.5 kPa were plotted as a function of the log_2_(fold change) in each comparison. Yellow boxes denote anti-correlated peaks. The top 2 transcription factor motifs enriched in the two anti-correlated peak sets identified by HOMER known motif analysis are shown. *P* values for motif analysis were calculated using HOMER (c-f). Pearson correlation coefficients (*R*) and *P* values were calculated using ggpubr in R via stat_cor() and regression line equations were estimated using stat_regline_equation() (e,f).

Differentially accessible peaks across sample comparisons were classified by genomic region. Matrix softening in wild-type WI-38 cells led to an increase of ATAC peaks in intronic and undefined regions of the genome (Fig. S6a). Additionally, mapping peaks to the nearest transcriptional start site identified that with substrate softening, peaks in wild-type cells mapped further from transcriptional start sites (Fig. S6b). These results were consistent with prior work^7^ demonstrating that accessibility increases in undefined genomic regions in senescent cells.

Generally, matrix softening led to an increase in chromatin accessibility (Fig. S6c). Motif analysis using HOMER^26^ of differentially accessible regions between conditions revealed striking patterns of transcription factor motif enrichment (Supplemental Data File 3). Matrix softening resulted in an increase in chromatin accessibility at activator protein-1 (AP-1) transcription factor component motifs (e.g. FOS, ATF3, JUN) specifically in wild-type WI-38 cells and not hTERT cells (Fig. 4c,d, S6d,e) (Supplemental Data File 3). Meanwhile, matrix stiffening as well as hTERT expression led to an increase in accessibility at CTCF-related motifs (Fig. 4c,d, S6d,e, Supplemental Data File 3).

If the increasing AP-1 motif accessibility with matrix softening in wild-type WI-38 cells contributed to their premature proliferative arrest, then these changes in accessibility would be expected to intensify over time. To determine the most robust accessibility changes across the 7 weeks, “anti-correlated” peaks were identified, defined as those whose accessibility increased over time in one condition while simultaneously decreasing in accessibility over time in a second condition. This method allowed us to pinpoint opposing processes most likely to contribute to the phenotypic divergence of wild-type cells on soft substrates. After identifying anti-correlated peaks between wild-type cells on stiff and soft substrates, motif enrichment analysis was performed. These analyses revealed that matrix softening resulted in an increase in AP-1 motif accessibility over time, in both promoter and genome-wide ATAC peaks of wild-type cells (Fig. 4e, S6f). Simultaneously, AP-1 motif accessibility decreased over time on stiff substrates (Fig. 4e, S6f). Additionally, comparison of wild-type and hTERT cells on soft substrates revealed AP-1 motif accessibility to inversely change between genotypes, increasing in wild-type cells and decreasing in hTERT-expressing cells across time (Fig. 4f, S6g).

The AP-1 transcription factor is a dimer that comprises proteins from the JUN, FOS, ATF, and/or MAF families^27,28^. AP-1 regulates diverse transcriptional programs in response to cellular stress, growth factors, cytokines, and more, and the composition of the AP-1 dimer determines the activation of specific gene expression programs^27,29^. To discern which AP-1 subfamilies may drive the increased motif accessibility in wild-type cells with matrix softening, motif sequences were compared to known binding elements of AP-1 family proteins. The AP-1 motifs identified by HOMER closely resembled 12-O-tetra-decanoylphorbol-13-acetate (TPA) response elements (TREs) rather than other motifs bound by AP-1 family members (Fig. 4c-f, S6d-h, Supplemental Data File 3)^27,30^. TREs are preferentially bound by JUN and FOS family proteins rather than ATF and MAF proteins, suggesting that matrix softening induced JUN and/or FOS DNA binding in wild-type WI-38 cells^27,28,30,31^.

### Environmental stiffness and hTERT status affect the formation of unique G1 cell states

ATAC-sequencing revealed the presence of distinct chromatin states dependent on matrix stiffness and genotype. To further investigate the formation of cellular states, scRNA-sequencing was performed on wild-type and hTERT cells after 1, 4, and 7 weeks of culturing on different matrices (Fig. 5a). Over 100,000 individual cells were profiled, and 9 cell clusters were identified after performing dimensionality reduction and clustering (Fig. 5b, S7a,b). Using established markers for S and G2/M phases, cells were classified by cell cycle phase^32^. Wild-type and hTERT cells across all matrix stiffnesses occupied the three S and G2/M phase clusters (clusters 0, 5, and 6), suggesting similarity in actively dividing cells across conditions (Fig. 5c-e, S7c-e).

**Figure 5:**
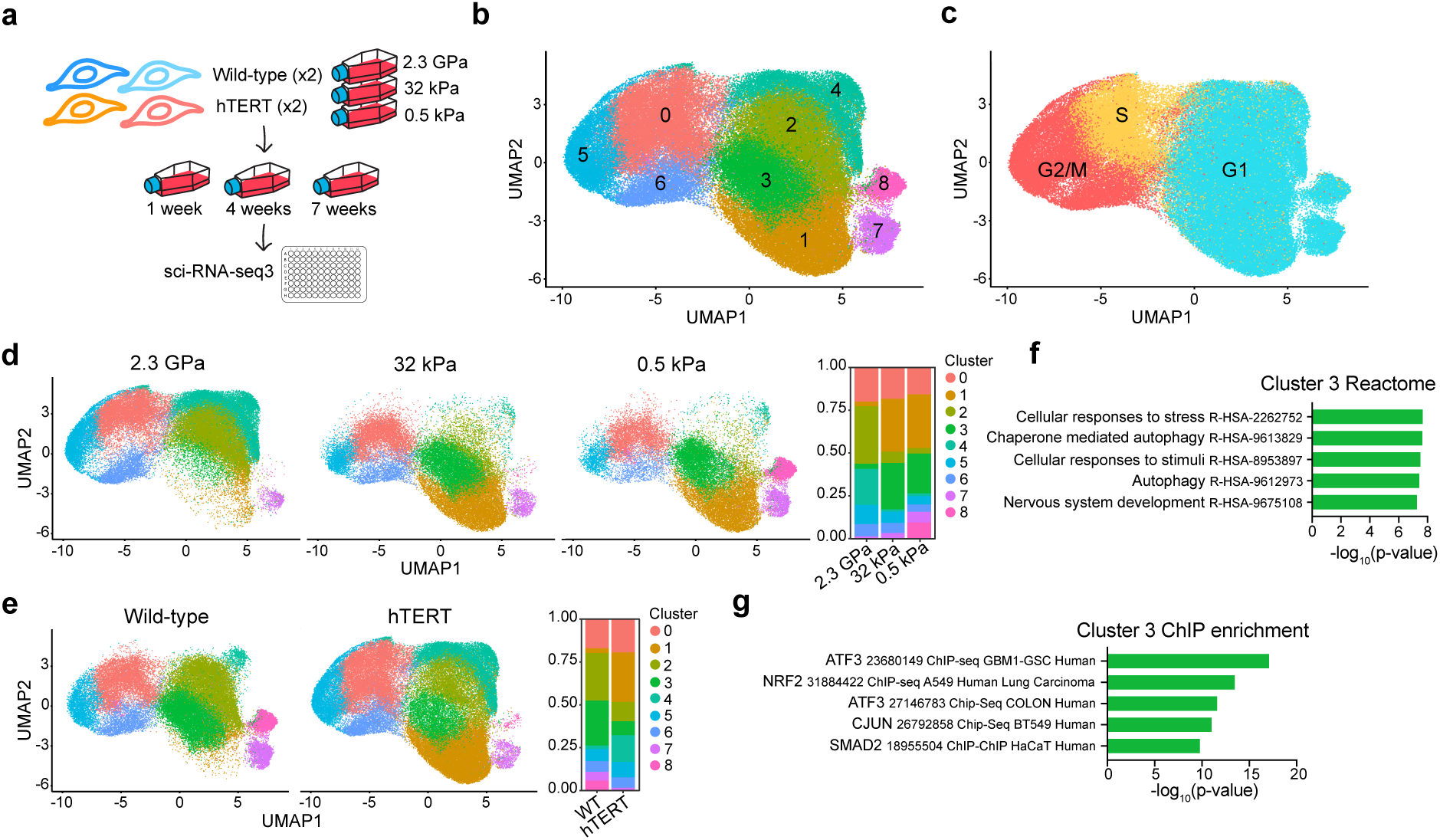
Formation of distinct G1 states depends on matrix stiffness and hTERT status. (a) Schematic for the scRNA-sequencing experiment. (b) UMAP projection showing UMAP1 and UMAP2 colored by cluster. (c) UMAP plot from (b) colored by different phases of the cell cycle. (d,e) (Left) UMAP plot from (b) split by (d) matrix stiffness or (e) genotype. (Right) Bar graph displaying the relative proportion of cells in each cluster. (f) Top 5 significantly enriched terms in cluster 3 markers in the 2022 Reactome pathway database via Enrichr^33–35^. (g) Top 5 significantly enriched chromatin immunoprecipitation (ChIP) datasets in cluster 3 markers using ChIP enrichment analysis via Enrichr^33–35^.

In contrast, G1 cluster formation was influenced by genotype and environmental stiffness (Fig. 5c-e, S7c-e). Cells expressing senescence-associated signatures appeared almost exclusively in wild-type cells on the softest 0.5 kPa substrate (Fig. 5d-e, S7f). For each of the most populated G1 clusters (clusters 1-4), the majority of cells derived from a unique combination of substrate stiffness and wild-type/hTERT status: wild-type cells on stiff or soft matrices in clusters 2 and 3, respectively, and hTERT cells on stiff or soft matrices in clusters 4 and 1, respectively (Fig. 5d-e, S7c). Leveraging this separation, markers of each cluster were used to identify pathways active in each population (Supplemental Data File 4). While the majority of markers were specific to each cluster, genes upregulated in clusters 1, 2, and 4 were associated with shared biological pathways, including ECM organization and collagen biosynthesis (Fig. S7g, Supplemental Data File 4). In contrast, markers of G1 cells from wild-type cells on soft matrices were uniquely associated with stress signaling pathways (Fig. 5f). To identify putative transcription factors that may mediate this transcriptional shift, chromatin immunoprecipitation (ChIP) enrichment analysis of cluster 3 markers was performed. G1 phase wild-type cells on soft substrates were characterized by an upregulation of genes bound by AP-1 transcription factor components (Fig. 5g). Thus, the enhanced AP-1 motif accessibility on soft matrices in wild-type WI-38 cells was concordant with the transcriptional profile of the distinct G1 population that emerged under those conditions.

### *JUNB* is upregulated as cell proliferation slows on soft matrices

ATAC- and scRNA-sequencing revealed a distinct opening of JUN/FOS binding motifs and activation of AP-1 target genes in G1 phase when wild-type WI-38 cells were moved to softer matrices. We revisited our bulk RNA-sequencing data to further explore the connection between JUN/FOS expression, matrix stiffness, and proliferative lifespan in WI-38 cells. The JUN family consists of the c-JUN (*JUN*), JUNB, and JUND proteins and the FOS family comprises c-FOS (*FOS*), FOSB, FOSL1, and FOSL2^27^. As differential expression of individual components can modulate dimer composition^27^, expression patterns of these seven genes were analyzed in the bulk RNA-sequencing dataset. All three JUN transcripts as well as *FOSL1* and *FOSL2* were expressed (Fig. S8a). However, *FOSL1* and *FOSL2* were more highly expressed in hTERT cells compared to wild-type cells as well as cells grown on stiff compared to soft matrices, suggesting these genes were likely not mediating the increase in AP-1 activity in wild-type cells with matrix softening (Fig. S8b,c).

Meanwhile, JUN family proteins’ transcripts were upregulated in wild-type cells relative to hTERT cells (Fig. S8c). *JUN* and *JUND* showed variable expression over time in wild-type cells across matrix types (Fig. 6a). *JUNB*, however, was steadily induced over time in wild-type cells on soft matrices (Fig. 6a). Similarly, when cells were grown on the intermediate stiffness matrices, *JUNB* expression was upregulated across time (Fig. S8d). On the stiff substrate, *JUNB* expression remained relatively constant until replicative senescence, when *JUNB* was induced (Fig. 6a). To understand how matrix stiffness regulated *JUNB*, its expression was examined when cells started to exhibit growth arrest in response to matrix softening (time point 4) (Fig. 2a). At this critical time, *JUNB*, and not *JUN* or *JUND*, showed upregulation with matrix softening (Fig. 6b). Concurrently, a number of known JUNB target genes^36–38^ demonstrated patterns of expression consistent with *JUNB* induction (Fig. S8e). For example, *CDKN2A*, which harbors the RNA transcript for *JUNB* target gene *p16^INK4A^*, was upregulated in wild-type cells on soft relative to stiff substrates (Fig. S8e). Together, the patterns of *JUNB* induction observed and previous work connecting JUNB with suppressing fibroblast proliferation (see Discussion) identified JUNB as a potential mediator of the premature WI-38 proliferative arrest with matrix softening.

**Figure 6:**
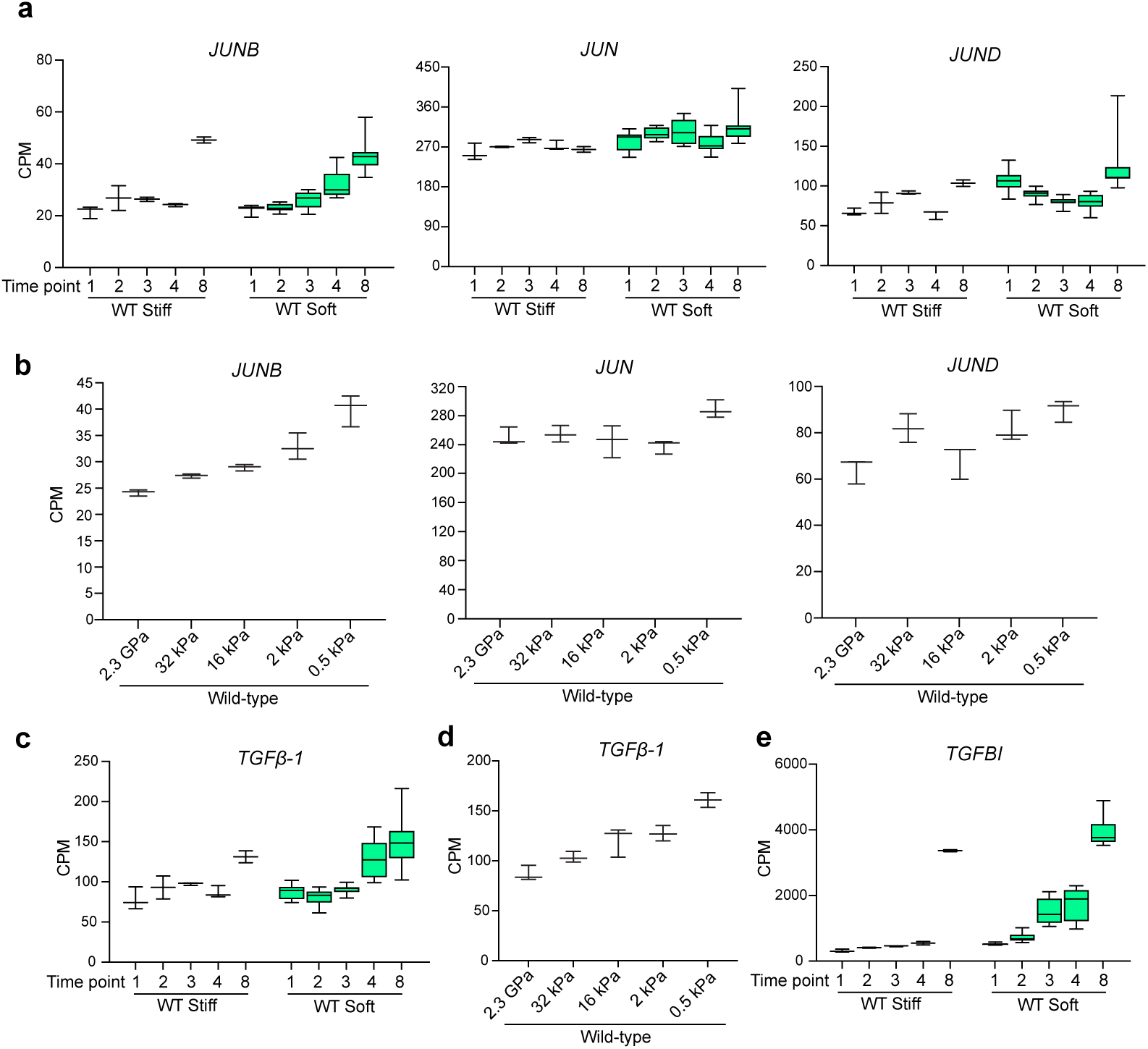
*JUNB* is upregulated as cells arrest with matrix softening. (a) Box plots of *JUN* family transcript expression across time in wild-type WI-38 cells in the initial time course RNA-sequencing experiment. (b) *JUN* family transcript expression at time point 4 in wild-type cells across substrate stiffnesses. (c) Box plots of *TGFβ-1* expression across time in wild-type cells. (d) *TGFβ-1* expression at time point 4 in wild-type cells across substrate stiffnesses. (e) Box plot of *TGFBI* expression across time in wild-type WI-38 cells. Box plots show median ± 25th and 75th percentiles. Whiskers show minimum to maximum.

Diverse mechanisms can induce *JUNB* transcription, including the MAPK pathway, TGF-β signaling, STAT3 activation, and reactive oxygen species^37,39–43^. Focusing on previously identified transcriptional activators of *JUNB* as candidate mediators of its expression, we identified several genes whose patterns of expression were similarly upregulated over time on soft matrices, specifically in wild-type cells. For example, TGF-β pathway signaling is known to induce *JUNB*^43^, and TGFβ1 expression and TGF-β target genes showed a time and matrix softening-dependent increase in expression (Fig. 6c-e, S8f,g). Similarly, the *JUNB* regulator STAT3^41,42^ and associated target genes were upregulated over time on soft matrices (Fig. S8h-k).

## Discussion

Studies have previously revealed intriguing links between aging phenotypes and the mechanical environment. For example, stiffening of the oligodendrocyte progenitor cell microenvironment with age is sufficient to impair progenitor functions, and growing aged chondrocytes on soft matrices restores gene expression programs reminiscent of younger cells^44,45^. Such studies, however, have been limited to investigating the short-term impacts of altering substrate stiffness. Long-term cellular responses to changes in environmental stiffness, including its effect on replicative lifespan, are not understood. Here, we performed the first longitudinal study of how substrate stiffness affects the biology and lifespan of WI-38 cells, a classic model of cellular aging. We discovered that manipulating matrix stiffness had extensive effects on WI-38 cell biology that were consistent across a wide range of elastic moduli and different substrates. Matrix softening induced a premature proliferative arrest, the development of discrete G1 cell states, and the induction of *JUNB* expression with concomitant chromatin opening at AP-1 transcription factor binding sites. Ultimately, these changes resulted in wild-type WI-38 cells undergoing fewer population doublings before undergoing growth arrest.

### Short-term impacts of substrate stiffness on gene expression did not persist over time

We hypothesized that moving cells to softer matrices would dampen the induction of YAP, EMT, and myofibroblast gene expression programs previously observed in WI-38 cells undergoing replicative senescence, potentially lengthening replicative lifespan^7^. Attenuation of these replicative senescence-associated programs was previously observed with acute verteporfin treatment^7^. However, WI-38 cells were intolerant to long-term verteporfin exposure at doses high enough to elicit a transcriptional response, driving our decision to alter mechanotransduction via environmental stiffness (Fig. S9a, data not shown). In the initial response to matrix softening, as expected, many of the YAP target genes previously associated with replicative senescence^7^ were downregulated; however, their levels rebounded and increased over time (Fig. S3b,f,g). Other YAP targets were initially upregulated by the softer mechanical environment and further increased in expression with replicative age (Fig. S3f,g). In all, the short-term changes in expression of YAP targets driven by environmental stiffness was not sufficient to overcome their dynamics in response to replicative age. These observations undermined our hypothesis, suggesting instead that replicative lifespan triggers the YAP signaling pathway orthogonally from mechanical cues.

Like YAP target genes, EMT genes also increase in expression when WI-38 cells approach replicative senescence on hard surfaces^7^. On soft substrates, EMT genes exhibited varied initial responses but ultimately increased through time (Fig. 2d, S3c). A similar pattern was again observed for myofibroblast markers that are associated with replicative senescence on stiff surfaces (Fig. S3d,e)^7^. These consistent patterns of short-term modification of gene expression by substrate stiffness, followed by long-term reversion to the expression patterns associated with replicative age, highlighted that paradigms of acute responses to substrate stiffness may not apply to long-term responses. These observations bolstered the conclusion that mechanical environments and replicative age may provide orthogonal stimuli for highly overlapping transcriptional programs.

### Expression of hTERT suppresses early indications of senescence-like phenotypes on soft substrates

Changes in gene expression previously established as specific to replicative senescence^7^ were broadly consistent as WI-38 cells approached proliferative arrest, regardless of substrate stiffness. Nonetheless, multiple cellular phenotypes varied substantially depending on the mechanical environment. First, the proliferation kinetics differed from cells undergoing replicative senescence: the cellular growth rate slowed earlier on softer substrates (Fig. S9b). Second, average telomere length was not critically shortened on soft matrices ahead of replicative arrest (Fig. 3e). Third, despite consistent behavior of senescence-relevant gene expression programs across stiffnesses, distinct epigenomic and transcriptional programs emerged with matrix softening that were also influenced by hTERT expression (Figs. 4,5). These observations indicated the existence of unique paths towards proliferative arrest dependent on environmental stiffness.

While matrix softening induced a proliferative arrest and morphological changes associated with senescence in wild-type WI-38 cells, hTERT-expressing cells, after an initial reduction in growth rate, showed no signs of a growth arrest after 5 months of culturing across matrix stiffnesses. hTERT expression rescued many morphological, proliferative, transcriptional, and epigenetic phenotypes induced by matrix softening. Previously, TERT has been shown to be insufficient to prevent various forms of premature stress-induced senescence, such as via ionizing radiation or oxidative damage^46,47^. Here, however, TERT rescued the premature growth arrest induced by matrix softening in the absence of critical telomere shortening.

Multiple mechanisms could explain the early influence of TERT on matrix softening-induced cellular phenotypes. One possibility is that the same DNA damage response to short telomeres on stiff substrates occurs earlier on soft matrices. This would require soft substrates to either increase the variance of telomere length (thereby allowing extreme degradation of one or more telomeres) or reduce the threshold of telomere shortening needed to trigger replicative senescence. In our study, while telomere loss per unit time was similarly rapid across substrates, we note that telomere loss per population doubling accelerated on soft substrates (Fig. S9d). However, when cellular phenotypes manifested, no accompanying transcriptional activation of DNA damage response markers was detected on soft matrices (Fig. S9c)^48–50^. An alternate mechanism by which TERT may influence matrix-softening associated phenotypes is through a non-canonical molecular function. TERT has been implicated in a myriad of pathways distinct from telomere lengthening, including regulating DNA methylation, the response to reactive oxygen species, NF-κB, and more^51^. Moreover, recent studies have identified that TERT regulation of DNA methylation modulates aging phenotypes^22^ and revealed a role of TERT in enhancing stem cell proliferation independent of telomere shortening^52^. Given how hTERT expression rescues proliferative and epigenetic phenotypes induced by matrix softening prior to drastic telomere shortening, the possibility that non-classical TERT functions and mechanosensation are intertwined becomes intriguing.

### JUNB as a novel transducer of mechanosensory cues

While the strong influence that the mechanical environment exerts on cellular behavior has been established in a variety of contexts^44,45,53,54^, the molecular mechanisms through which mechanosensory signals influence the phenotypes associated with proliferative arrest, as we observed here, are unknown. Moreover, TERT suppression of mechanoresponsive phenotypes was unexpected and has no known mechanistic basis. In order to connect these phenomena, we sought to identify transcription factors whose patterns of expression and activity implicated them as potential mediators.

Enrichment analysis for transcription factor binding motifs was performed in DNA regions that either gained accessibility when wild-type cells were grown on soft versus stiff substrates or lost accessibility on soft substrates when expressing TERT. In both cases, these analyses revealed motifs bound by AP-1 transcription factor complexes, and scRNA-sequencing confirmed the importance of AP-1 regulated genes in defining distinct transcriptomic states in response to those perturbations. AP-1 proteins are broadly considered as pro-proliferative and oncogenic^27^ – seemingly contrary to the proliferative arrest induced with matrix softening. However, context and dimer composition dictate functions across the diverse AP-1 family. The single G/C nucleotide in between two palindromic TGA’s in our detected motifs identified them as being of the TRE sub-class of AP-1 motifs, known to be bound by JUN homodimers or JUN/FOS heterodimers^27^. Among those candidates, upregulation of *JUNB* particularly correlated with the emergence of soft substrate-induced phenotypes and transcriptomic signatures, implicating JUNB as a potentially key mediator of those phenomena.

Several known aspects of JUNB biology support a central role in the long-term phenotypic responses of WI-38 cells to soft substrates. JUNB can mediate anti-proliferative effects in fibroblasts, at least partially through induction of the cyclin-dependent kinase inhibitor *p16^INK4A^* ^36,39^. Given the established anti-proliferative role of JUNB, the upregulation of *JUNB* with matrix softening over time, and chromatin opening at TRE elements, we hypothesize that AP-1 dimers increasingly comprise JUNB proteins over time with matrix softening and initiate anti-proliferative transcriptional programs. The question of how *JUNB* is activated with matrix softening remains, but our data supported several candidates. Multiple known inducers of *JUNB* expression were upregulated on soft matrices over time, including *TGFβ-1* and *STAT3*^37,41,42^. While the TGF-β and STAT pathways have been linked to mechanosensation, their upregulation with matrix softening has not been reported^55–57^. Additionally, how TERT might interact with this pathway to rescue proliferation and other phenotypes associated with soft matrices remains to be explored.

By growing WI-38 cells on soft substrates for extended periods of time, we were able to characterize novel phenotypic evolutions as cells age. In this long-term growth context, the JUNB component of the AP-1 transcription factor was implicated as a key mediator. With JUNB not having been previously associated with mechanosensation, our findings illustrate the importance of longitudinal culturing of cells on different matrices, identifying new time- and stiffness-dependent patterns of gene expression. This study also revealed an unexpected interaction between TERT expression and the mechanoresponsive behavior of cells. Both telomere shortening and tissue stiffening are hallmarks of the aging process. Our observation of TERT modifying the responses of cells to shifting mechanical environments reveals a novel mechanism that increases the interconnectedness of aging phenotypes, which in turn may contribute to the exponentially accelerating physiological decay that accompanies human aging.

## Methods

### Cell culture

WI-38 cells (Coriell) and hTERT WI-38 cells (generated in-house, see below) were maintained in low glucose Dulbecco’s Modified Eagle medium (DMEM) (Gibco) supplemented with 10% fetal bovine serum (Gibco) and 1% Antibiotic-Antimycotic (Gibco) in an incubator set to 37°C, 20% O_2_, and 5% CO_2_. Cells were routinely tested for mycoplasma and always tested negative. Cells were grown on plastic (polystyrene) plates (Falcon), CytoSoft (PDMS) plates (Advanced Biomatrix), polyacrylamide gels (4Dcell), or PDMS plates fabricated in-house. All plates were coated with PureCol Type I collagen (Advanced Biomatrix). To collagen-coat plates, PureCol was diluted 1:40 in water (for plastic plates) or 1x phosphate buffered saline (PBS, Corning) (for PDMS and polyacrylamide gels) and incubated on the plate for 2 hours at room temperature. Plates were washed 2x with PBS and then tissue culture media was added. When cells reached 70-80% confluence, cells were passaged by washing the plate with PBS and then incubating with TrypLE Express Enzyme (Gibco) until cells lifted from the plate. Media was added to neutralize the reaction and the cell suspension was spun at 300 x *g* for 5 minutes. The resulting cell pellet was resuspended in media and counted using a ViCell XR Cell Analyzer with Trypan Blue staining to assess cell viability (Beckman Coulter). Each sample was counted twice, and the resulting counts were used for population doubling calculations. Population doubling level was calculated as follows:

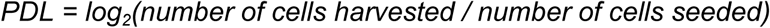

For verteporfin experiments, verteporfin (R&D Systems) was diluted in DMSO. Cells were seeded into plates and treated with the vehicle control (DMSO) or verteporfin (0.001, 0.01, 0.1 1, or 10 μm) every 72 hours.

### Lentivirus generation and transduction

293FT cells (Invitrogen) were grown in sterile-filtered DMEM (high glucose, no glutamine, no sodium pyruvate) (Gibco) supplemented with 10 mM MEM non-essential amino acids (Gibco), 100 mM MEM sodium pyruvate (Gibco), and 200 mM GlutaMax (final concentration: 6mM) (Gibco). Cells were transfected with a lentiviral plasmid containing hTERT (pCDH-CMV-hTERT-EF1a-puro) and lentiviral packaging constructs using Lipofectamine 2000 (ThermoFisher). WI-38 cells were infected with viral supernatant containing 5 µg/mL polybrene (Fisher) and selected for 1 week with 1 µg/mL puromycin (Sigma). hTERT expression was confirmed by quantitative polymerase chain reaction.

### Senescence-associated β-galactosidase staining

Wild-type and hTERT WI-38 cells were periodically split into 6-well collagen-coated plastic or CytoSoft plates for SA-β-gal staining. Cells were stained using the Senescence β-Galactosidase Staining Kit (Cell Signaling) following the manufacturer’s instructions. At least 2 wells and 100 cells were counted for each condition.

### PDMS plate fabrication

To fabricate polydimethylsiloxane (PDMS) plates of variable stiffnesses, PDMS and curing agent (RTV615) (R.S. Hughes) were mixed at different ratios (Supplemental Data File 1). Materials were mixed vigorously and then spun and deaerated using an ARE-310 THINKY mixer. The resulting mixture was pipetted into plastic tissue culture dishes (Falcon) and degassed via vacuum. Plates were then incubated in an oven set to 80°C for variable amounts of time to achieve different stiffnesses (Supplemental Data File 1). The day before use, PDMS gels underwent plasma cleaning and silane treatment to reduce hydrophobicity. Plates were placed in a plasma cleaner (Harrick Plasma) and underwent plasma cleaning for 1 minute. A 10% silane (Sigma) in ethanol solution was prepared and added to gels for 15 minutes with frequent agitation. Plates were washed 5x with 100% ethanol and then 4x with 70% ethanol. Plates were placed in an 80°C oven to dry for 1 hour. When cooled, plates were collagen coated as described above. To measure the elastic moduli of the gels, samples were sent to EAG Laboratories for nanoindentation analysis.

### RNA-sequencing

For all bulk RNA-sequencing experiments, cells were lysed directly on cell culture plates in Buffer RLT Plus (Qiagen). The lysate was homogenized using Qiashredder columns (Qiagen) and stored at -80°C. Once all samples were collected, RNA was isolated using an RNeasy mini kit (Qiagen). Quality and concentration of RNA were determined using a 2100 Bioanalyzer Instrument (Agilent). cDNA libraries were constructed for the initial time course (comprising stiff and soft matrices) using the NEBNext Ultra II Directional RNA Library Prep Kit for Illumina with the rRNA depletion workflow (New England Biolabs). cDNA libraries were constructed for the intermediate stiffness time course (comprising stiff, intermediate, and soft matrices) using the NEBNext Ultra II Directional RNA Library Prep Kit for Illumina with the polyA mRNA workflow (New England Biolabs). Samples were sequenced using a NovaSeq6000 (Illumina). FASTQ files were aligned to the human genome (hg38) and sorted by genomic location using STAR (v.2.5.3, https://github.com/alexdobin/STAR). Sorted BAM files were counted and mapped to each gene using HTSeq-Count (v.0.11.3, https://github.com/simon-anders/htseq). Differentially expressed genes (DEGs) were identified using DESeq2 (v.1.44.0, https://github.com/thelovelab/DESeq2)^58^ in R (v.4.2.2) with a significance cutoff of *P*-adjusted < 0.05. Wald test was used to calculate fold change and *P* values using the model: ∼condition, where condition is a variable capturing time point, genotype, and substrate stiffness. Principal component analysis and sample-to-sample distance mapping were performed using DESeq2. Heatmap visualization was performed using pheatmap (v.1.0.12, https://github.com/raivokolde/pheatmap). The EMT, YAP targets, and myofibroblast signatures visualized in heatmaps were previously identified^7^, and the EMT signature is listed in Supplemental Data File 2.

### Telomere qPCR

For telomere qPCR experiments, cells were harvested from plates using TrypLE Express Enzyme as described above and stored frozen in liquid nitrogen in Recovery Cell Culture Freezing Medium (Gibco). Once all samples were collected, telomere qPCR was performed as previously described^23–25^. Briefly, DNA was isolated using the Monarch Genomic DNA Purification Kit (New England Biolabs). DNA concentration was measured in triplicate on a NanoDrop One (Thermo Scientific). Samples were diluted to 5 ng/µl, and qPCR was performed in triplicate using 20 ng of DNA per reaction with PowerUp SYBR Green Master Mix (Applied Biosystems) and primers targeting telomeres (forward primer: 5’ CGGTTTGTTTGGGTTTGGGTTTGGGTTTGGGTTTGGGTT 3’; reverse primer: 5’ GGCTTGCCTTACCCTTACCCTTACCCTTACCCTTACCCT 3’) and an internal single-copy reference gene control, *36B4* (forward primer: 5’ CAGCAAGTGGGAAGGTGTAATCC 3’; reverse primer: 5’ CCCATTCTATCATCAACGGGTACAA 3’). qPCR was performed using a QuantStudio 6 Real-Time PCR System (Applied Biosystems). Cycling conditions: 10 minutes at 95°C, followed by 40 cycles of 95°C for 15 seconds, 60°C for 1 minute. Results were analyzed using the 2^-ΔΔCT^ method.

### ATAC-sequencing

For the ATAC-sequencing experiment, cells were harvested from plates using TrypLE Express Enzyme as described above and stored frozen in liquid nitrogen in Recovery Cell Culture Freezing Medium (Gibco). Once all samples were collected, ATAC-sequencing was performed as previously described^59^. In brief, frozen cells were thawed rapidly in a 37°C water bath and 7.5x10^4^ cells were aliquoted and washed once with PBS. Nuclei were isolated, lysed, and incubated with Tn5 transposase (Illumina) as described. Transposed DNA was isolated with the MinElute Reaction Cleanup Kit (Qiagen). Libraries were prepared by amplifying DNA for 5 cycles with NEBNext 2x MasterMix (New England Biolabs). After performing qPCR to determine additional cycles needed, libraries were amplified for an additional 6 cycles. Double sided bead purification was performed using AMPure XP beads (Beckman) to remove primers and large (>1000 bp) fragments using standard protocols. Quality and concentration of libraries were determined using a Qubit 4 Fluorometer (Invitrogen) and a 2100 Bioanalyzer Instrument (Agilent). Samples were sequenced using a NovaSeq6000 (Illumina). FASTQ files were trimmed and low quality reads were filtered using cutadapt (v.4.9.0, https://github.com/marcelm/cutadapt) to remove primer sequences.

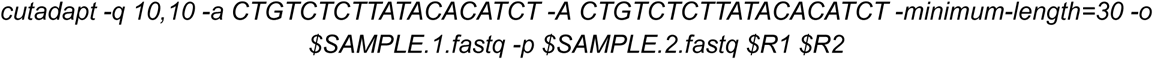

Trimmed FASTQ files were then aligned to the genome using bowtie2 (v.2.3.4, https://github.com/BenLangmead/bowtie2), and sorted based on genomic location and indexed using Samtools (v.1.14.0, https://github.com/samtools/samtools). Quality control, peak calling, differential peak analysis, and principal component analyses were performed using ChrAccR (v.0.9.17, https://github.com/GreenleafLab/ChrAccR) in R (v.4.3.0). Default parameters were used to generate the promoter peaks list. To generate the genome-wide peaks list (all peaks) via MACS2 (v.2.2.9.1, https://pypi.org/project/MACS2), the setConfigElement doPeakCalling() was used with annotationPeakGroupAgreePerc() = 1. PAVIS^60^ was performed for peak annotation with the upstream and downstream length parameters set to 5 kb (https://manticore.hiehs.nih.gov/pavis2). The genomic regions enrichment of annotation tools (GREAT, v.4.0.4, https://great.stanford.edu) was used to identify the distance of peaks from transcriptional start sites. Transcription factor motif analyses were performed individually on differentially accessible peaks from each comparison using the HOMER motif discovery tool (v.4.11.0, https://homer.ucsd.edu/homer) with the findMotifsGenome() command.

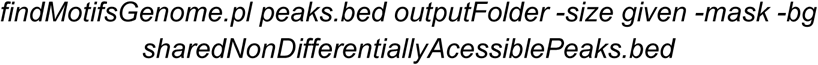

Anti-correlated peaks were identified by identifying all differentially accessible regions over time (time 1 versus time 7) in two conditions (e.g. wild-type cells on the 0.5 kPa substrate versus wild-type cells on the stiff substrate). After plotting the log_2_(fold change) over time in the two conditions against each other, anti-correlated peaks were identified as those showing opposite directionality of the log2(fold change) in each condition. Pearson correlation coefficients and significance were calculated using ggpubr (v.0.6.0, https://github.com/kassambara/ggpubr) via stat_cor() and regression line equations were estimated using stat_regline_equation().

### Single-cell RNA-sequencing

For the scRNA-sequencing experiment, cells were harvested from plates using TrypLE Express Enzyme as described above and stored frozen in liquid nitrogen in Recovery Cell Culture Freezing Medium (Gibco). Once all samples were collected, single-cell combinatorial indexing RNA-sequencing (sci-RNA-seq3) was performed as previously described^61^. In brief, the plate-based single-cell RNA-sequencing method from Scale Biosciences was used for processing the samples. Cryopreserved cells were thawed in a water bath and recovered in media (10% FBS in 1x low glucose DMEM). The cells were further washed 2 times in 1x PBS and resuspended in 1x PBS for fixation. The fixed samples were stored in -80°C for 1 week and processed through the ScaleBio single-cell RNA sequencing kit (v1.1). The library was sequenced on a NovaSeq6000. The resulting data was processed through bcl2fastq and further processed through the ScaleRna pipeline (v1.4.0, https://github.com/ScaleBio/ScaleRna) to generate the gene-by-cell count matrices. The resulting expression matrices were loaded into R (v.4.3.0) and processed using Seurat (v4.3.0, https://github.com/satijalab/seurat) for downstream analyses. Cells were filtered to only include cells with over 200 RNA features, less than 11,000 RNA features, and less than 5% mitochondrial DNA reads. SCTransform() was performed to normalize, scale, find variable features, and regress the variable number of RNA counts. Principal component analysis was run to identify the first 50 major axes of variation using RunPCA(). Batch integration was then performed using Harmony (v.1.0.0, https://portals.broadinstitute.org/harmony). The variable used in group.by.vars was the batch day samples from the same time point were harvested.

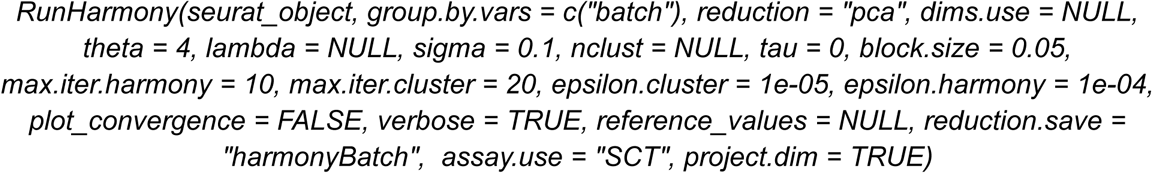

Clustering was performed by constructing a nearest neighbor graph using FindNeighbors() and identifying clusters of cells by a shared nearest neighbor modularity optimization-based clustering algorithm using FindClusters(). Clusters were visualized by UMAP dimensionality reduction via RunUMAP(). The following parameters were used to generate the final UMAP: # of PCs = 12, k = 30, resolution = 0.4. FindAllMarkers() was used to identify specific gene markers of each cluster. Cell cycle scoring was performed using the Seurat implementation of the cell cycle scoring function, as previously described^32^. Enrichr (v.3.2.0, https://maayanlab.cloud/Enrichr)^33–35^ was used to perform gene ontology and ChIP enrichment analyses. Gene module scores of senescence programs were calculated by AddModuleScore(). Pearson correlation coefficients of quality control data were calculated in R using Seurat via the FeatureScatter() function.

### Statistical analysis

Data analysis and statistical tests were performed using GraphPad Prism software (v.10.2.0). For the ATAC-sequencing analysis, Pearson correlation coefficients and related *P* values as well as regression line equations were computed using ggpubr (v.0.6.0, https://github.com/kassambara/ggpubr) in R. For the scRNA-sequencing analysis, Pearson correlation coefficients were calculated using Seurat (v4.3.0, https://github.com/satijalab/seurat). Student’s *t*-tests were two-tailed. All statistical tests are denoted in the figure legends.

## Data availability

Data from the bulk RNA-sequencing experiments is available at the GEO under the accession numbers GSE276045 and GSE276046. Data from the ATAC-sequencing experiment is available at the GEO under the accession number GSE276047. Data from the scRNA-sequencing experiment is available at the GEO under the accession number GSE276048.

## Acknowledgements

We thank D. Hendrickson and the Calico Genomics lab for their guidance on and processing of high-throughput sequencing samples.

## Contributions

A.M.K designed and performed experiments, interpreted and analyzed data, and wrote the manuscript. A.S. developed the methodology to fabricate tunable stiffness PDMS plates. W.K. performed the sci-RNA-seq3 and single-cell data processing up through the ScaleRNA pipeline. J.G.R. designed and interpreted experiments and wrote the manuscript. All authors reviewed and approved the final manuscript for submission.

## Supplemental files

**Supplemental Figures:** Nine Supplemental Figures (S1-S9) with legends as one PDF.

**Supplemental Data File 1:** Nanoindentation analysis results (Excel notebook file).

**Supplemental Data File 2:** Senescence-associated gene expression programs used in single cell cluster definition (Excel notebook file).

**Supplemental Data File 3:** HOMER analysis results (Excel notebook file).

**Supplemental Data File 4:** Markers of single cell clusters (Excel notebook file).

