## Supplemental Figures for "Substrate stiffness dictates unique paths towards proliferative arrest in WI-38 cells"

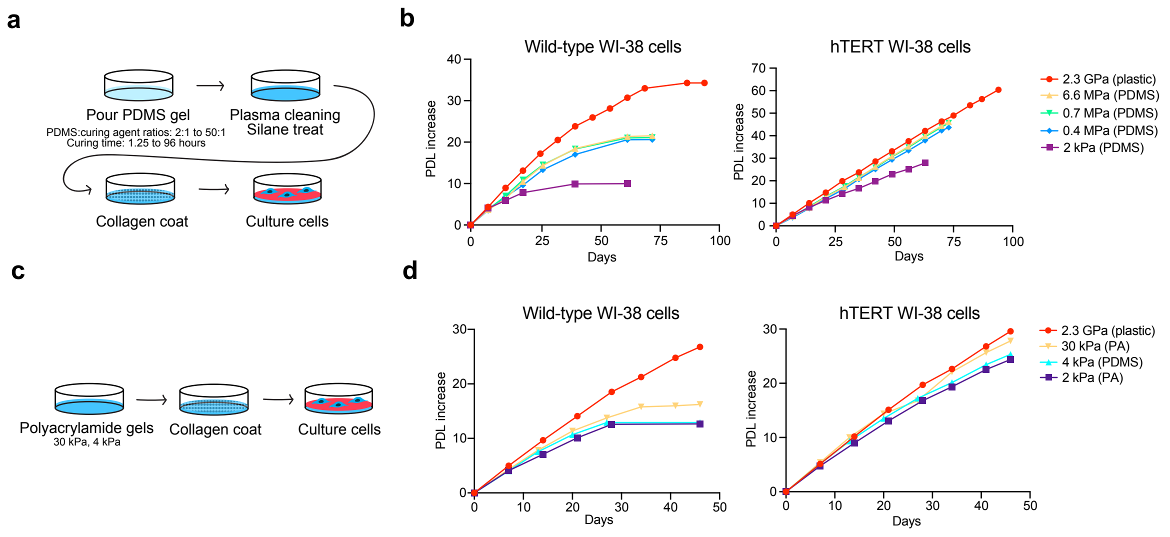


**Figure S1: Wild-type WI-38 cells show impaired proliferation across multiple types of soft substrates.**

(a) Schematic depicting PDMS plate fabrication. (b) Line graph displaying days in culture (x-axis) versus the PDL increase from day 0 (y-axis) for (left) wild-type or (right) hTERT WI-38 cells on the indicated matrices. (c) Schematic for polyacrylamide gel experiments. (d) Line graph displaying days in culture (x-axis) versus the PDL increase from day 0 (y-axis) for (left) wild-type or (right) hTERT WI-38 cells on the indicated matrices.

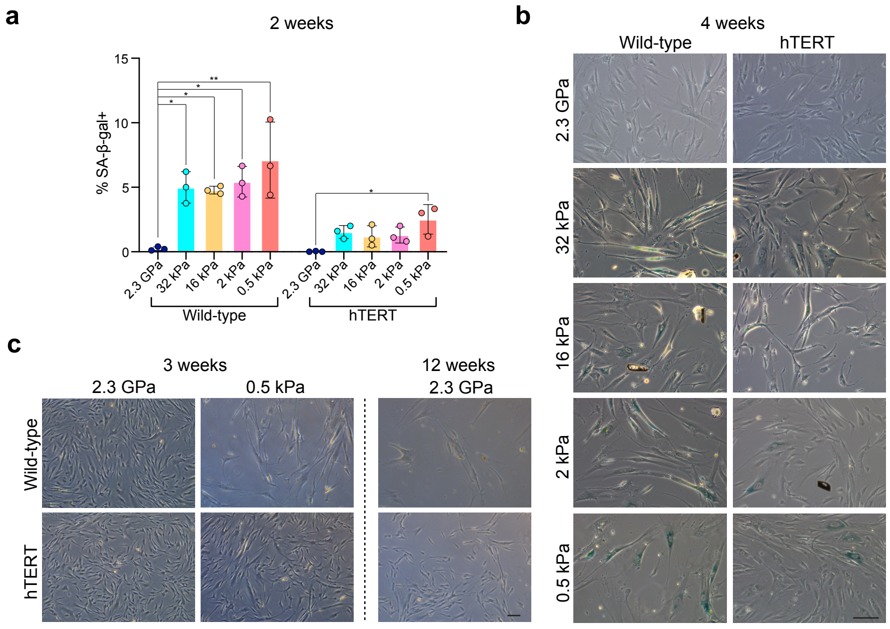


**Figure S2: Wild-type WI-38 cells on soft matrices exhibit classic features of senescent cells.**

(a) Bar graphs quantifying the percent of cells that stained for SA-β-gal in wild-type and hTERT-expressing WI-38 cells after 2 weeks in culture. (b) Representative images of SA-β-gal staining after 4 weeks of culture. (c) Representative images of wild-type and hTERT cells on the 2.3 GPa and 0.5 kPa surfaces at the indicated time points.

Bar graphs are mean ± standard deviation. *P* values were calculated by ordinary one-way ANOVA with Tukey's multiple comparisons test. $^*P < 0.05$, $^{**}P < 0.01$. Scale bars, 100 μm.

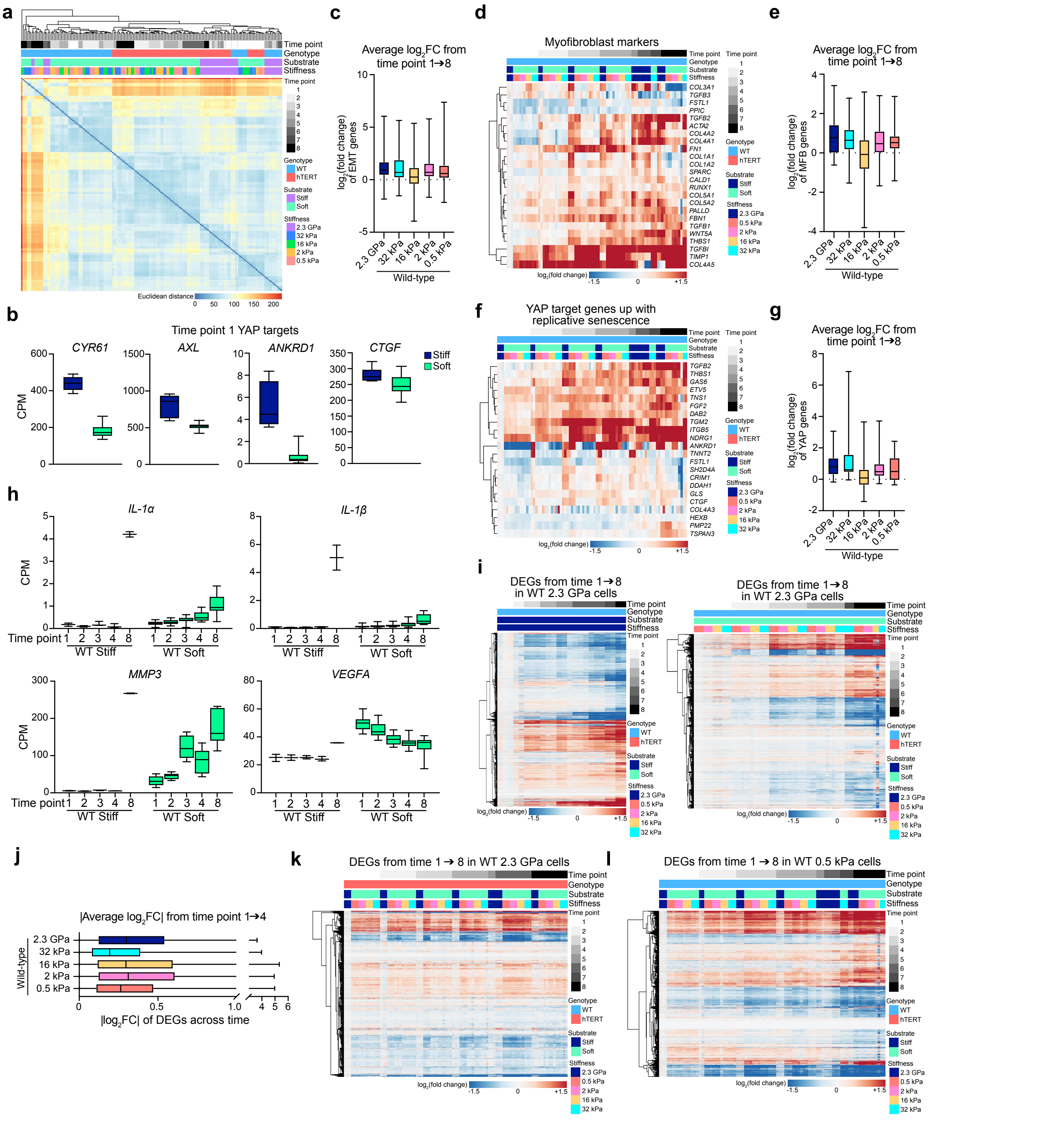


**Figure S3: WI-38 cell gene expression programs are similar across substrate stiffnesses.** (a) Heatmap of the Euclidean distance between RNA-sequencing samples. (b) Box plots comparing the expression of YAP target genes in WI-38 cells on stiff and soft matrices at time point 1. (c) Box plots quantifying the $\log_2$(fold change) of each gene in (Fig. 2d) from time point 1 to time point 8 averaged across replicates. (d) Heatmap of the $\log_2$(fold change) of myofibroblast (MFB) markers previously identified[7] as upregulated in replicative senescence. (e) Box plots quantifying the $log_2$(fold change) of each gene in (d) from time point 1 to time point 8 averaged across replicates. (f) Heatmap of the $log_2$(fold change) of YAP target genes previously identified[7] as upregulated in replicative senescence. (g) Box plots quantifying the $log_2$(fold change) of each gene in (f) from time point 1 to time point 8 averaged across replicates. (h) Box plots comparing the expression of SASP genes[22] in wild-type WI-38 cells on the indicated surfaces across time. (i) Heatmap from (Fig. 2f) split by wild-type cells grown on (left) stiff or (right) soft substrates. (j) Box plots quantifying the absolute value of the $log_2$(fold change) of each gene in (Fig. 2f) from time point 1 to time point 4 averaged across replicates. (k) Heatmap of the $log_2$(fold change) of genes from (Fig. 2f) in hTERT samples across time. (l) Heatmap of the $log_2$(fold change) of all DEGs in wild-type cells grown on the 0.5 kPa matrix from time point 1 to time point 8 plotted across all wild-type samples across time.

Box plots show median ± 25th and 75th percentiles. Whiskers show minimum to maximum.

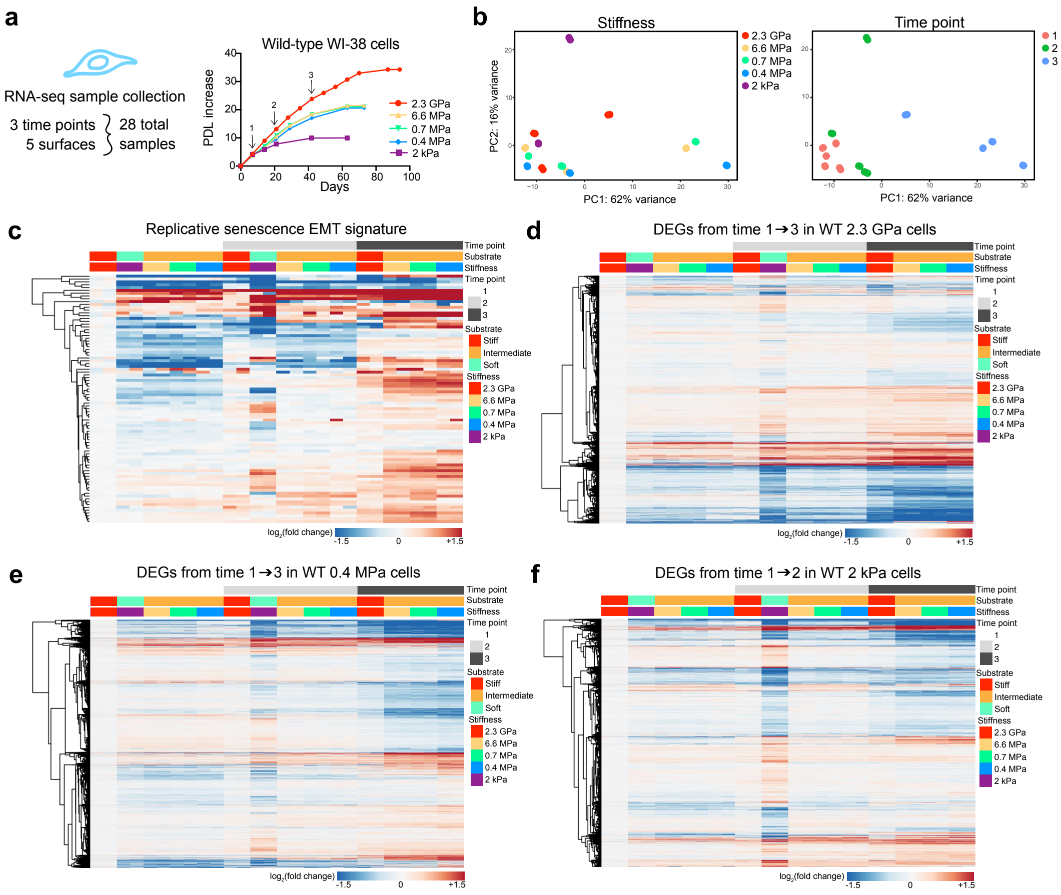


**Figure S4: WI-38 cells cultured on a wide range of PDMS stiffnesses similarly regulate senescence-associated gene expression programs.**

(a) Schematic of the (left) RNA-sequencing samples and (right) sample collection time points. (b) PC analysis showing PC1 and PC2 with samples colored by different variables. (c) Heatmap of the $\log_2$(fold change) of the EMT gene set (Supplemental Data File 2) previously identified[7] as upregulated in replicative senescence. (d-f) Heatmap of the $\log_2$(fold change) of all DEGs in wild-type cells grown on (d) 2.3 GPa, (e) 0.4 MPa, or (f) 2 kPa from time point 1 to time point 2 or 3 plotted across all samples across time.

**a**

| Sample ID | Sample identity | Sample ID | Sample identity | Sample ID | Sample identity |
|---|---|---|---|---|---|
| 1 | WT1 2.3GPa time 1 | 13 | WT1 2.3GPa time 4 | 25 | WT1 2.3GPa time 7 |
| 2 | WT2 2.3GPa time 1 | 14 | WT2 2.3GPa time 4 | 26 | WT2 2.3GPa time 7 |
| 3 | hTERT1 2.3GPa time 1 | 15 | hTERT1 2.3GPa time 4 | 27 | hTERT1 2.3GPa time 7 |
| 4 | hTERT2 2.3GPa time 1 | 16 | hTERT2 2.3GPa time 4 | 28 | hTERT2 2.3GPa time 7 |
| 5 | WT1 0.5kPa time 1 | 17 | WT1 0.5kPa time 4 | 29 | WT1 0.5kPa time 7 |
| 6 | WT2 0.5kPa time 1 | 18 | WT2 0.5kPa time 4 | 30 | WT2 0.5kPa time 7 |
| 7 | hTERT1 0.5kPa time 1 | 19 | hTERT1 0.5kPa time 4 | 31 | hTERT1 0.5kPa time 7 |
| 8 | hTERT2 0.5kPa time 1 | 20 | hTERT2 0.5kPa time 4 | 32 | hTERT2 0.5kPa time 7 |
| 9 | WT1 32kPa time 1 | 21 | WT1 32kPa time 4 | 33 | WT1 32kPa time 7 |
| 10 | WT2 32kPa time 1 | 22 | WT2 32kPa time 4 | 34 | WT2 32kPa time 7 |
| 11 | hTERT1 32kPa time 1 | 23 | hTERT1 32kPa time 4 | 35 | hTERT1 32kPa time 7 |
| 12 | hTERT2 32kPa time 1 | 24 | hTERT2 32kPa time 4 | 36 | hTERT2 32kPa time 7 |

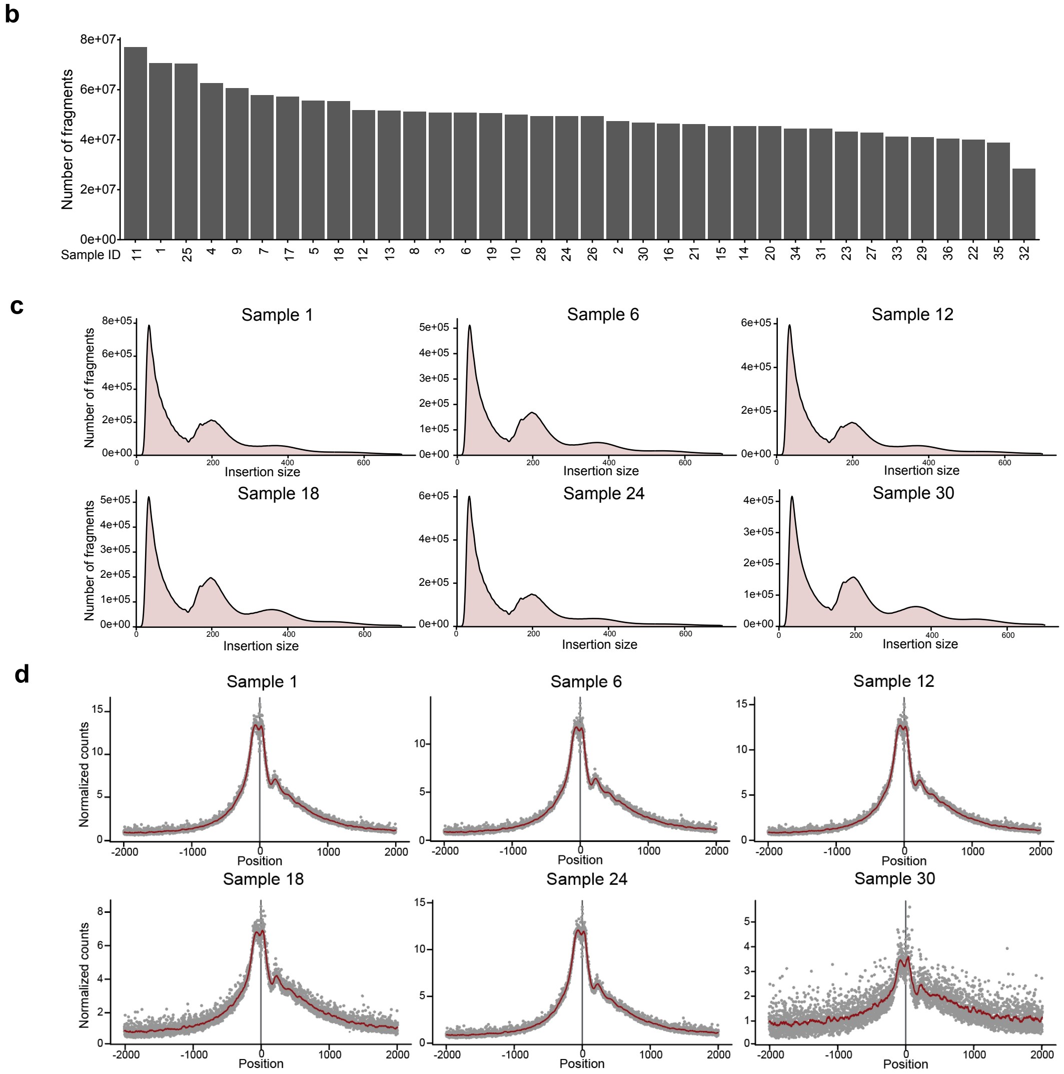


**Figure S5: ATAC-sequencing library quality controls.**

(a) Chart depicting the identity of ATAC-sequencing samples. (b) Bar graph displaying the fragment count of each ATAC-sequencing sample. (c) Fragment size distribution graphs of 6 different ATAC libraries shows a periodicity indicative of nucleosome bound DNA. (d) Plot of normalized counts per ATAC peak in the region ± 2 kilobases from the transcriptional start site to show enrichment of ATAC signal near the transcriptional start site of 6 different ATAC libraries.

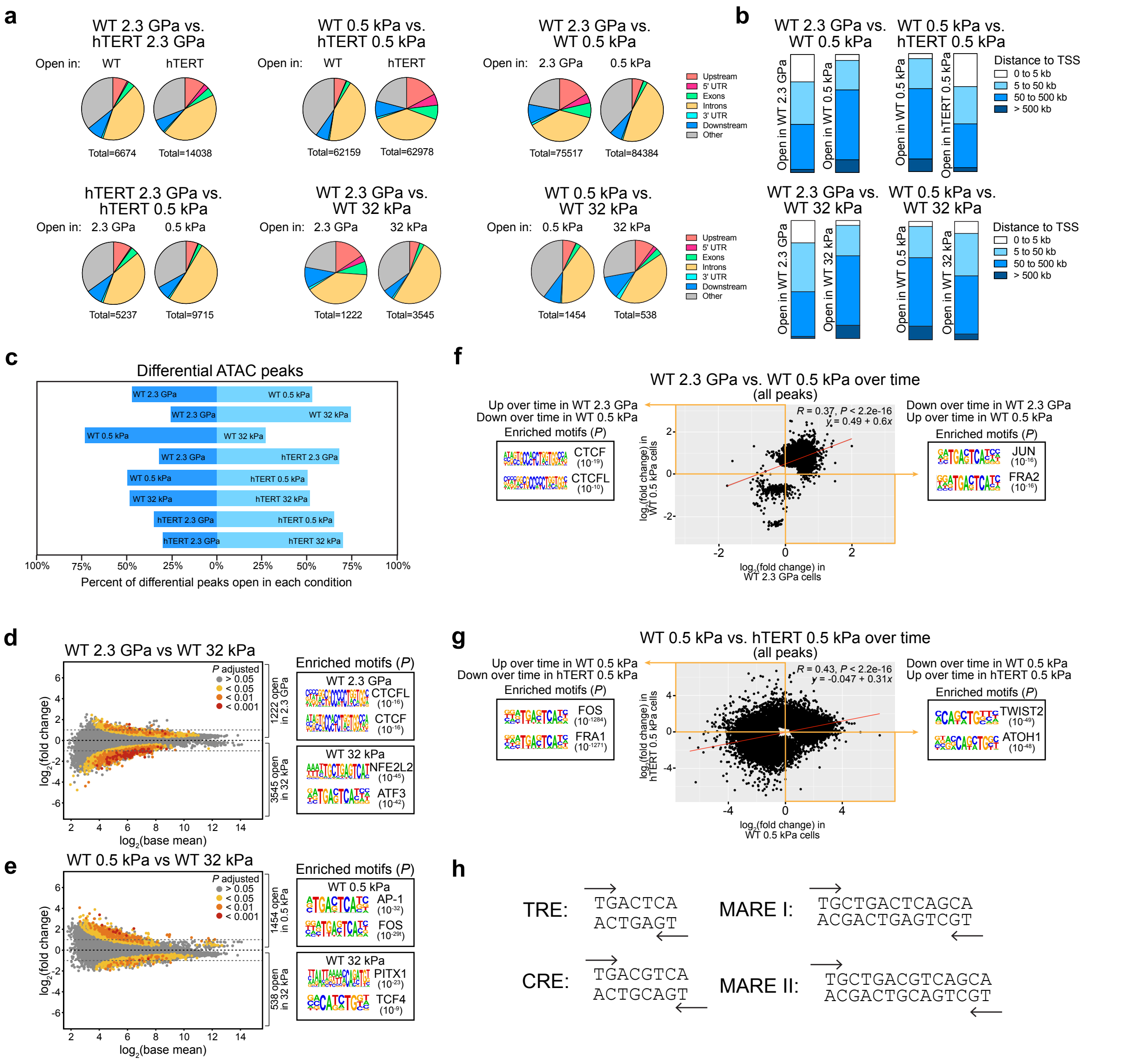


**Figure S6: Patterns of chromatin accessibility change with matrix stiffness.**

(a) Peak annotation of differentially accessible (DA) regions in the stated comparisons. UTR, untranslated region. (b) Distance of DA regions in the stated comparisons from the nearest transcriptional start site (TSS), plotted as a proportion of all DA regions. (c) Proportion of DA peaks (all peaks analysis) opening on each side of the stated comparisons. (d,e) $Log_2$ mean reads per region versus differential accessibility [$log_2$(fold change)] for peaks in the (d) wild-type 2.3 GPa versus wild-type 32 kPa comparison or (e) wild-type 0.5 kPa versus wild-type 32 kPa comparison. The top 2 transcription factor motifs enriched in DA regions identified by HOMER known motif analysis are shown. (f,g) All DA regions over time (time 1 versus time 7) in (f) wild-type cells on 2.3 GPa versus wild-type cells on 0.5 kPa or (g) wild-type cells on 0.5 kPa versus hTERT cells on 0.5 kPa were plotted as a function of the $log_2$(fold change) in each comparison. Yellow boxes highlight anti-correlated peaks. The top 2 transcription factor motifs enriched in the two anti-correlated peak sets identified by HOMER known motif analysis are shown. (h) TPA response element (TRE), cAMP-response element (CRE), and MAF-recognition element (MAF) sequences[7,27].

*P* values for motif analysis were calculated using HOMER (d-g). Pearson correlation coefficients (*R*) and *P* values were calculated using ggpubr in R via stat_cor() and regression line equations were estimated using stat_regline_equation() (f,g).

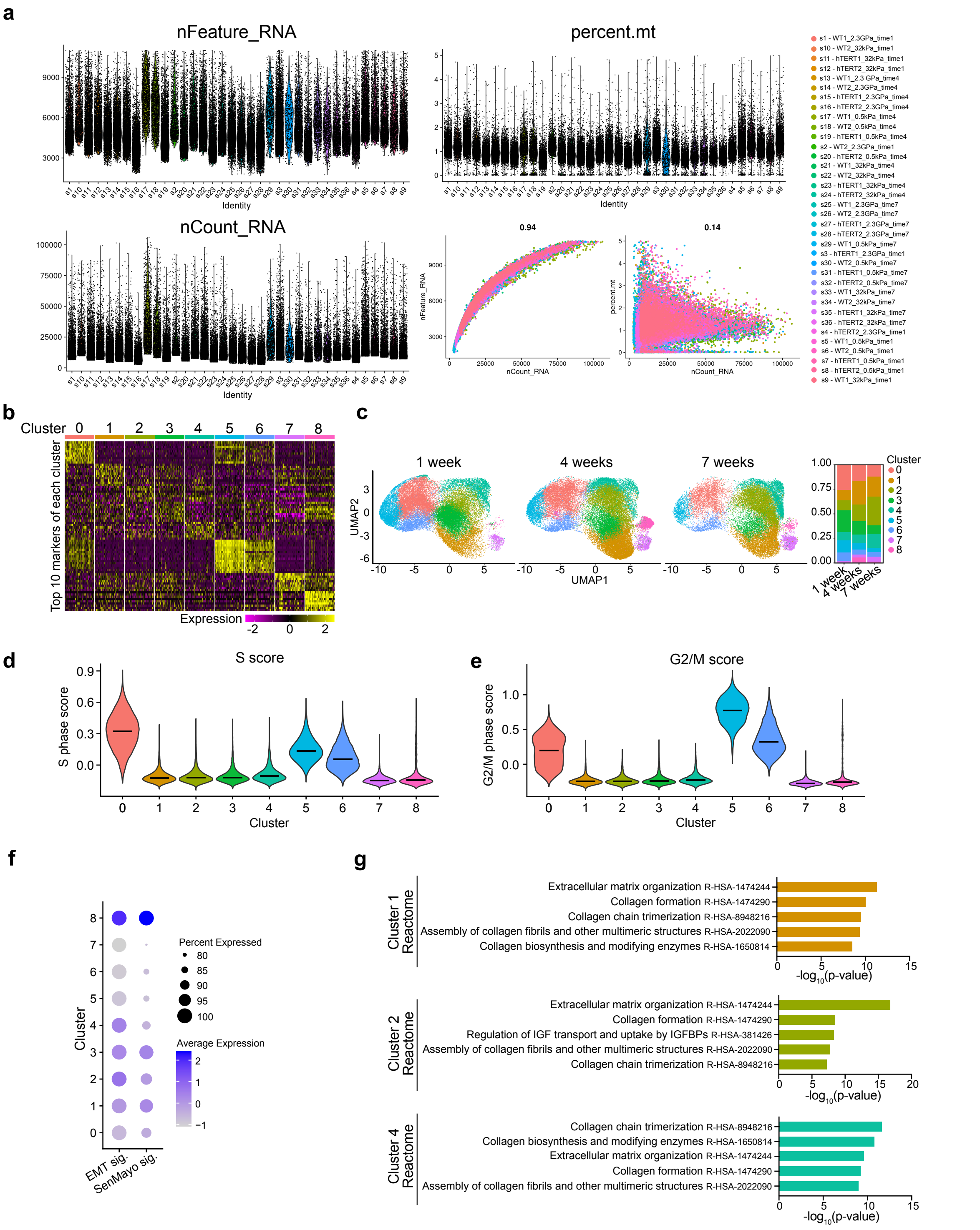


**Figure S7: scRNA-sequencing reveals distinct cell clusters.**

(a) Violin plots depicting the number of RNA features (nFeature_RNA), percent mitochondrial (mt)DNA (percent.mt), and the number of RNA counts (nCount_RNA) for individual cells in each scRNA-sequencing sample after filtering. Scatter plots show (left) nCount_RNA versus nFeature_RNA for individual cells and (right) nCount_RNA versus percent.mt for individual cells.

(b) Heatmap of the top 10 markers for each cluster calculated in R using Seurat via the FindAllMarkers() function. (c) (Left) UMAP plot from (Fig. 5b) split by time point. (Right) Bar graph displaying the relative proportion of cells in each cluster split by time point. (d,e) Violin plots showing the distribution of (d) S phase and (e) G2/M phase cell cycle scoring across clusters calculated in R using Seurat via the CellCycleScoring() function. (f) Dot plot showing the average expression of two senescence-associated gene expression programs (Supplemental Data File 2) across clusters. (g) Top 5 significantly enriched terms in cluster 1, 2, or 4 markers in the 2022 Reactome pathway database via Enrichr[33–35].
Pearson correlation coefficients (a) were calculated in R using Seurat via the FeatureScatter() function. Violin plots show the median.

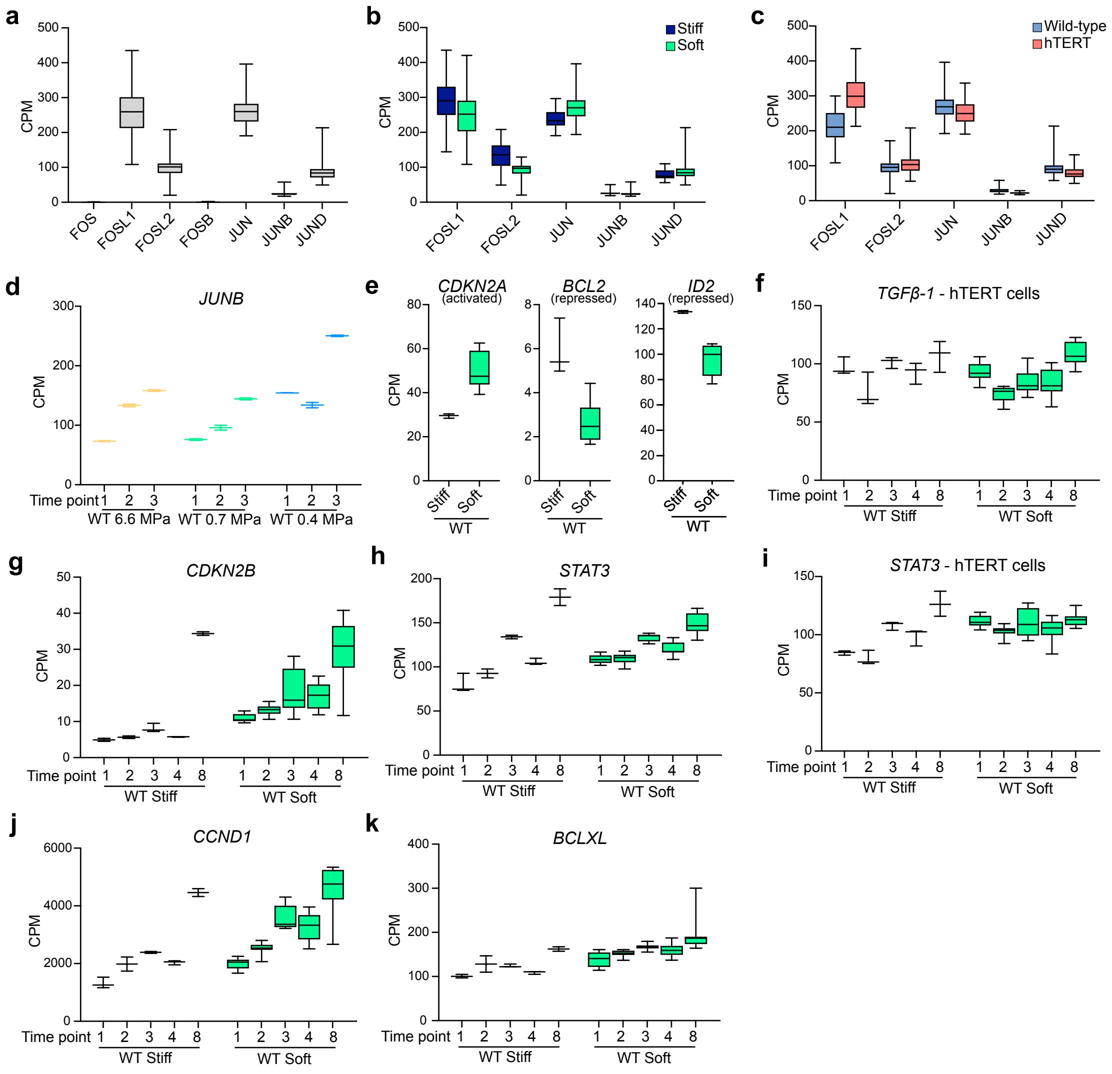

**Figure S8: *JUNB* and *JUNB* regulators are induced with matrix softening.**
(a-c) Box plots comparing the expression of JUN and FOS family members across all samples in the initial time course RNA-sequencing experiment. (d) *JUNB* expression across time in wild-type cells grown on substrates of intermediate stiffnesses. (e) Expression of JUNB target genes in wild-type cells at time point 4. (f) Box plots of *TGFβ-1* expression in hTERT cells. (g,h) Box plots of the indicated genes across time in wild-type cells. (i) Box plots of *STAT3* expression in hTERT cells. (j,k) Box plots of the indicated genes across time in wild-type cells.
Box plots show median ± 25th and 75th percentiles. Whiskers show minimum to maximum.

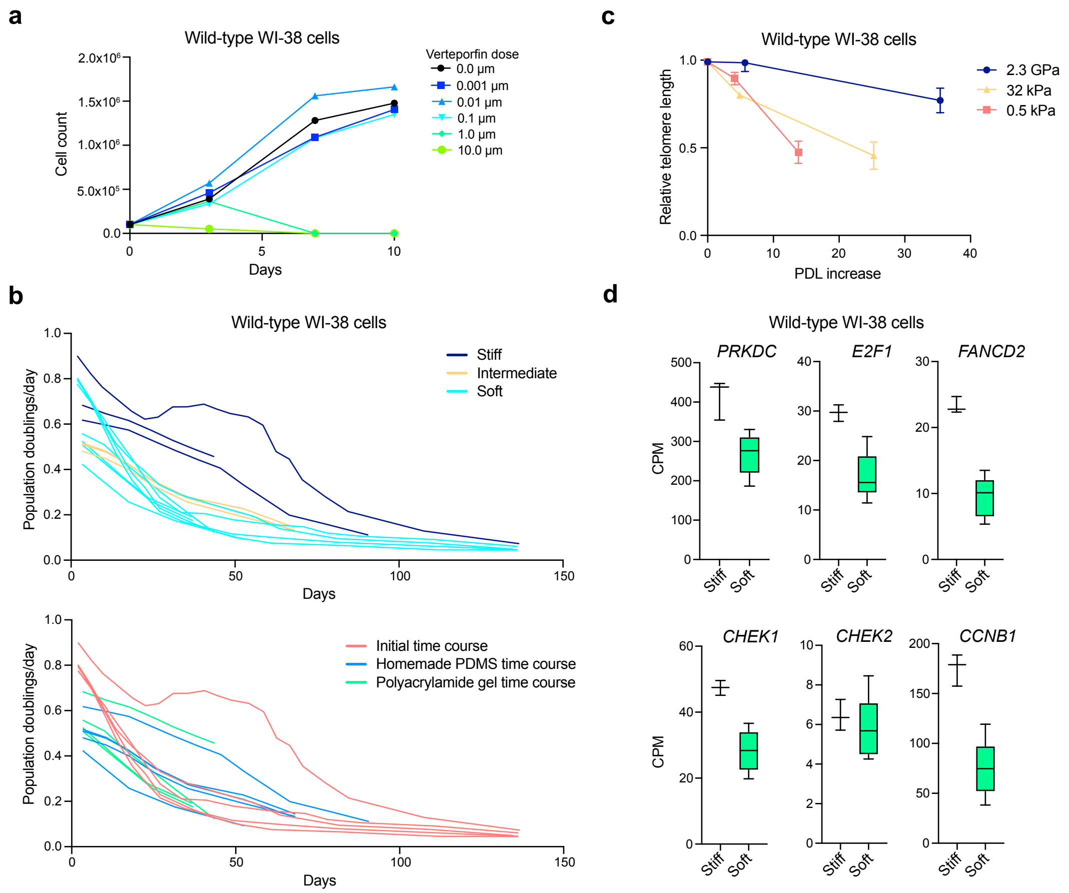


**Figure S9: Wild-type WI-38 growth rate and telomere degradation depend on substrate stiffness.**

(a) Line graph displaying days in culture (x-axis) versus the total number of wild-type WI-38 cells (y-axis) when cells were treated with different concentrations of verteporfin. (b) Line graphs displaying days in culture (x-axis) versus the number of population doublings per day (y-axis) underwent by wild-type WI-38 cells over time across all experiments colored by (top) substrate stiffness or (bottom) experiment. (c) Line graph displaying the PDL increase (x-axis) versus the relative telomere length (y-axis) for wild-type WI-38 cells on the indicated surfaces. (d) Expression of DNA damage response markers at time point 4 in wild-type WI-38 cells on the indicated stiffnesses in the initial time course RNA-sequencing experiment.

Box plots show median ± 25th and 75th percentiles. Whiskers show minimum to maximum.
